# *mlh3* separation of function and endonuclease defective mutants display an unexpected effect on meiotic recombination outcomes

**DOI:** 10.1101/108498

**Authors:** Najla Al-Sweel, Vandana Raghavan, Abhishek Dutta, V. P. Ajith, Luigi Di Vietro, Nabila Khondakar, Carol M. Manhart, Jennifer A. Surtees, K. T. Nishant, Eric Alani

**Affiliations:** Department of Molecular Biology and Genetics, Cornell University, Ithaca, NY, USA; School of Biology, Indian Institute of Science Education and Research Thiruvananthapuram, Trivandrum 695016, India; Department of Life Sciences and Systems Biology, University of Turin, Via Verdi, 8-10124, Turin, Italy.; Department of Biochemistry, University at Buffalo, State University of New York, Buffalo, NY, USA

**Author notes:** Present address: Bayer CropScience, Lyon, France (Luigi Di Vietro). Corresponding Author: Eric Alani, Cornell University, Department of Molecular Biology and Genetics, 459 Biotechnology Building, Ithaca, NY 14853-2703.

**Keywords:** Mlh1-Mlh3, meiotic crossing over, DNA mismatch repair, separation of function

## Abstract

Mlh1-Mlh3 is an endonuclease hypothesized to act in meiosis to resolve double Holliday junctions into crossovers. It also plays a minor role in eukaryotic DNA mismatch repair (MMR). To understand how Mlh1-Mlh3 functions in both meiosis and MMR, we analyzed in baker’s yeast 60 new *mlh3* alleles. Five alleles specifically disrupted MMR, whereas one (*mlh3-32*) specifically disrupted meiotic crossing over. Mlh1-mlh3 representatives for each separation of function class were purified and characterized. Both Mlh1-mlh3-32 (MMR^+^, crossover^-^) and Mlh1-mlh3-45 (MMR^-^, crossover^+^) displayed wild-type endonuclease activities *in vitro*. Msh2-Msh3, an MSH complex that acts with Mlh1-Mlh3 in MMR, stimulated the endonuclease activity of Mlh1-mlh3-32 but not Mlh1-mlh3-45, suggesting that Mlh1-mlh3-45 is defective in MSH interactions. Whole genome recombination maps were constructed for two *mlh3* mutants with opposite separation of function phenotypes, and an endonuclease defective mutant. Unexpectedly, all three showed increases in the number of non-crossover events that were not observed in *mlh3Δ*. Our observations provide a structure-function map for Mlh3 that reveals the importance of protein-protein interactions in regulating Mlh1-Mlh3’s enzymatic activity. They also illustrate how defective meiotic components can alter the fate of meiotic recombination intermediates, providing new insights for how meiotic recombination pathways are regulated.

**Author Summary:** During meiosis, diploid germ cells that become eggs or sperm undergo a single round of DNA replication followed by two consecutive chromosomal divisions. The segregation of chromosomes at the first meiotic division is dependent in most organisms on at least one genetic exchange, or crossover event, between chromosome homologs. Homologs that do not receive a crossover frequently undergo non-disjunction at the first meiotic division, yielding gametes that lack chromosomes or contain additional copies. Such events have been linked to human disease and infertility. Recent studies suggest that the Mlh1-Mlh3 complex is an endonuclease that resolves recombination intermediates into crossovers. Interestingly, this complex also acts as a matchmaker in DNA mismatch repair (MMR) to remove DNA replication errors. How does one complex act in two different processes? We investigated this question by performing a mutational analysis of the baker’s yeast Mlh3 protein. Five mutations were identified that disrupted MMR but not crossing over, and one mutation disrupted crossing over while maintaining MMR. Using a combination of biochemical and genetic analyses to further characterize these mutants we illustrate the importance of protein-protein interactions for Mlh1-Mlh3’s activity. Importantly, we illustrate how defective meiotic components can alter the outcome of meiotic recombination events. They also provide new insights in our understanding of the basis of infertility syndromes.

## Introduction

During mismatch repair (MMR), insertion/deletion and base-base mismatches that form as the result of DNA replication errors are recognized by <u>M</u>ut<u>S</u> <u>h</u>omolog (MSH) proteins, which in turn recruit <u>M</u>ut<u>L</u> <u>h</u>omolog (MLH) proteins to form ternary complexes containing mismatched DNA, MSH factors, and MLH factors. These interactions result in the recruitment of downstream excision and resynthesis proteins to remove the error [1]. In *S. cerevisiae* repair of insertion deletion loops greater than one nucleotide in size primarily involves the MSH heterodimer Msh2-Msh3 and the MLH heterodimer Mlh1-Pms1 [1]. The MLH heterodimer Mlh1-Mlh3, has been shown to play a minor role in this process and can partially substitute for Mlh1-Pms1 in Msh2-Msh3-dependent MMR [2-4]. However, Mlh1-Mlh3 has been shown to play a major role in meiotic crossing over [5-8]. Accurate chromosome segregation in Meiosis I in most eukaryotes requires reciprocal exchange of genetic information (crossing over) between homologs [9-12]. Failure to achieve at least one crossover (CO) per homolog pair results in homolog nondisjunction and the formation of aneuploid gametes. Errors in meiotic chromosome segregation are a leading cause of spontaneous miscarriages and birth defects [13].

Yeast Mlh1-Pms1 and its human ortholog MLH1-PMS2 both exhibit an endonuclease activity that is essential for MMR [14-15]. This activity is dependent on the integrity of a highly conserved (DQHA(X)_2_E(X)_4_E) metal binding motif also found in Mlh3. Previous work demonstrated that a point mutation within this motif (*mlh3-D523N*) conferred *mlh3Δ*-like defects in MMR and crossing over. These included a mutator phenotype, a decrease in spore viability to 70% (from 97% in wild-type), and a two-fold reduction in genetic map distances [5]. Consistent with these observations, Mlh1-Mlh3 is an endonuclease that nicks circular duplex DNA and whose activity is enhanced by Msh2-Msh3 *in vitro* [16-17].

Approximately 200 double strand breaks (DSBs) are induced throughout the genome in a *S. cerevisiae* cell in meiotic prophase, of which ~90 are repaired to form COs between homologous chromosomes, with the rest repaired to form non-crossovers (NCOs; [18-23]). In this pathway a DSB, which primarily forms on one chromatid of a homologous pair, is resected by 5’ to 3’ exonucleases, resulting in the formation of 3′ single-strand tails on both sides of the DSB (Fig 1). One of these tails invades the other unbroken homolog and is extended and stabilized to create a single-end invasion intermediate (SEI). A second invasion event initiating from the SEI, known as second-end capture, can re-anneal and ligate to the other side of the DSB resulting in the formation of a double Holliday junction (dHJ). The dHJ can be acted upon by Holliday junction (HJ) resolvases to form CO and NCO products. In baker’s yeast the majority of COs are formed through an interference-dependent CO pathway (class I COs) in which the vast majority of dHJs are resolved to form evenly spaced COs in steps requiring the ZMM proteins Zip1-4, Mer3, and Msh4-Msh5 as well as the Sgs1-Top3-Rmi1 (STR) helicase/topoisomerase complex, Mlh1-Mlh3, and Exo1 [8, 24-31]. These steps are biased to resolve the two junctions present in the dHJ in opposing orientations such that the resulting product is primarily a CO. A second interference-independent pathway was identified that accounts for a small (~10%) number of CO events (class II COs). In this pathway, which does not involve the ZMM proteins, the two junctions are resolved independently by the Mms4-Mus81 endonuclease, leading to a mixture of CO and NCO products [7, 8, 32, 33].

**Fig 1.**
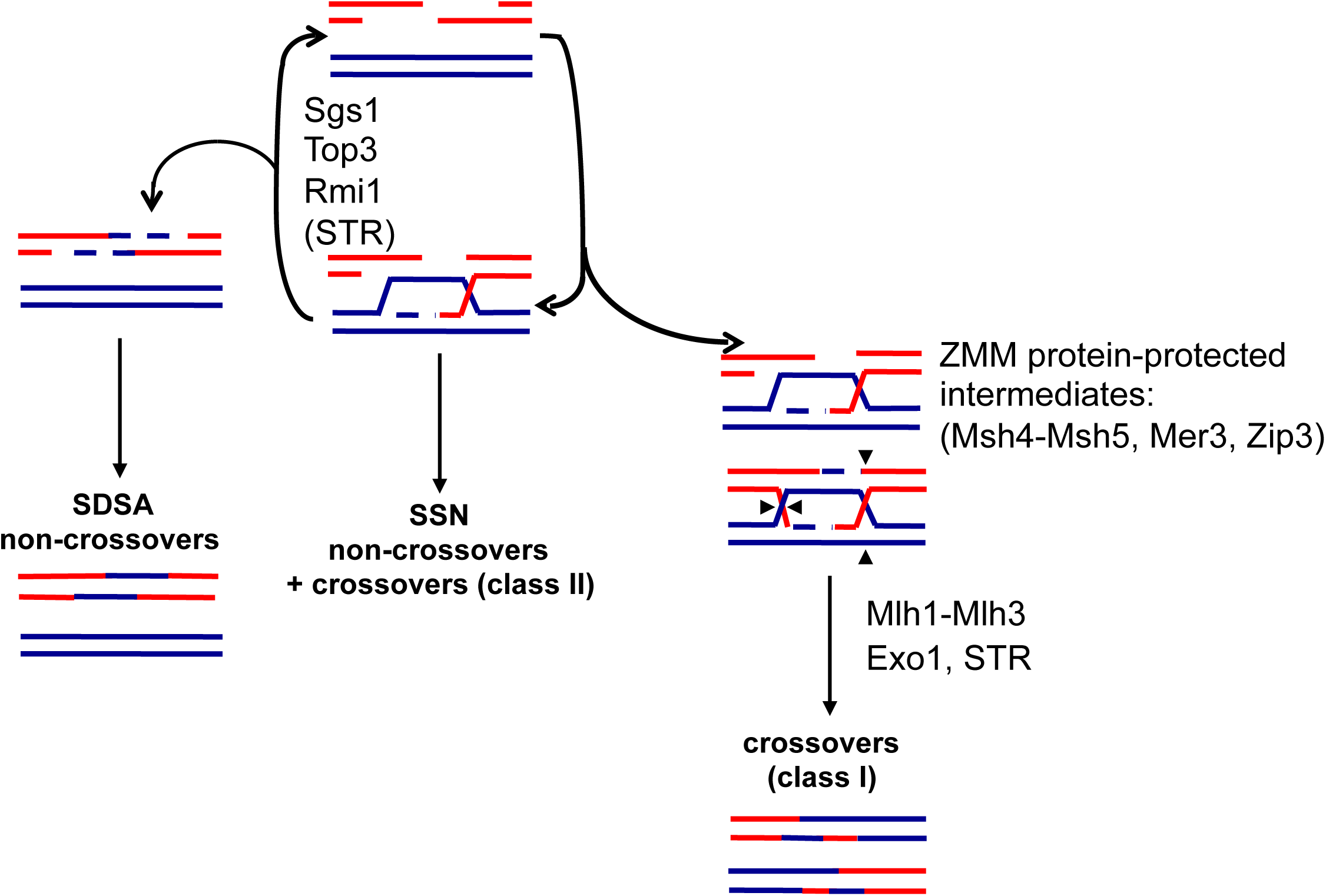
DSB repair pathways in meiosis. Model from Kaur et al. [30] depicting wild-type meiosis and the central role of the STR complex (Sgs1-Top3-Rmi1 helicase/topoisomerase) in disassembling strand invasion intermediates to facilitate synthesis dependent strand annealing (SDSA) or return of events to the original DSB state to allow capture and protection by the ZMM proteins and dHJ formation for ultimate resolution as class I crossovers by Mlh1-Mlh3 and Exo1. Events that escape STR disassembly form unregulated joint molecules that are resolved by the structure selective nucleases (SSNs) as noncrossovers or class II crossovers.

Genetic and physical studies, summarized below, support a major role for Mlh1-Mlh3 in promoting meiotic CO formation in the interference-dependent CO pathway. 1. Genetic studies showed that *mlh1* and *mlh3* mutants display approximately two-fold reductions in crossing over [7, 34, 35]. 2. There is significant redundancy of factors required to resolve dHJs into COs. This redundancy involves the endonucleases Mlh1-Mlh3, Mus81-Mms4, Yen1, and Slx1-Slx4 [5, 7, 8, 36], with Yen1 and Slx1-Slx4 acting in cryptic or backup roles. When all four factors were removed, crossing over was reduced to nearly background levels; however in an *mms4 slx4 yen1* triple mutant, in which Mlh1-Mlh3 is maintained, relatively high CO levels (~70% of wild-type levels) were observed, suggesting that Mlh1-Mlh3 is the primary factor required for CO resolution in the interference-dependent CO pathway [8]. 3. MLH1 and MLH3 play critical roles in mammalian meiosis [37, 38]. For example, *Mlh3*^*-/-*^ mice are sterile with an 85-94% reduction in the number of COs; germ cells in these mice fail to maintain homologous pairing at metaphase and undergo apoptosis [37, 39].

Much remains to be understood on how biased resolution of dHJs in the interference-dependent pathway is achieved. A working model, supported by genetic and molecular studies outlined below, is that the STR complex and a subset of ZMM proteins process and interact with DSB repair and SEI intermediates to create a dHJ substrate that can be resolved by the Mlh1-Mlh3 endonuclease and Exo1 to form primarily COs [5, 7, 8, 16, 29, 30, 31, 35, 36, 40-46]. In this model, the biased cleavage of a dHJ suggests coordination between the two junctions that would likely require asymmetric loading of meiotic protein complexes at each junction. However, little is known at the mechanistic level about how such coordination could be accomplished. A recent bioinformatics study by the Fung group, which involved the analysis of CO-associated gene conversion patterns in yeast tetrads, suggested that Zip3, a SUMO E3 ligase, is required for biased cleavage [47]. Curiously, they found that biased resolution of dHJs was maintained in *msh4* mutants. Based on these findings and other observations they propose that Msh4-Msh5 is required at the invading end of the DSB to stabilize recombination intermediates such as SEIs, while Zip3 acts to promote second-end capture steps at the ligating end of the DSB [47]. In support of this model, the ZMM heterodimer Msh4-Msh5 has been shown to promote COs in the same pathway as Mlh1-Mlh3, and human MSH4-MSH5 was shown to bind to SEI and Holliday junction substrates *in vitro* [43]. Furthermore, cytological observations in mouse have shown that MSH4-MSH5 foci appear prior to MLH1-MLH3 [37, 44, 48, 49]. Consistent with these observations, MLH1 and MLH3 foci formation requires MSH4-MSH5 [49].

Additional support for the above model was obtained from analysis of the STR complex, which has been identified as a pro-CO factor in the ZMM pathway [8, 30, 31, 46, 50]. The STR complex has recently been labeled the master regulator of meiotic DSB repair, acting as both a positive and negative CO coordinator (Fig 1) [30, 50]. Initially, the Sgs1 helicase was characterized as anti-CO because it facilitates unwinding of DSB repair intermediates. However, deleting either Sgs1 or Mlh3 in yeast strains that lack all other meiotic resolvases (*mms4, slx4, yen1*) results in a similar reduction of CO levels (~10% of wild-type levels) suggesting a pathway where Sgs1-dependent COs require Mlh1-Mlh3. Similar results were observed in *mms4, slx4, yen1* strains deficient in Top3 or Rmi1 [30, 31]. These data indicate that the STR complex promotes the majority of COs in conjunction with a resolvase that is not Mus81-Mms4, Slx1-Slx4 or Yen1.

A role for Exo1 in crossing over is supported by genetic studies that show Exo1 and Mlh3 acting in the same CO pathway [29]. Interestingly, Exo1’s role in maintaining wild-type levels of crossing over is independent of its catalytic activity, suggesting a structural role for this pro-CO factor [29]. Consistent with the above observations, Msh4-Msh5, STR, Exo1 and Zip3 have all been shown to interact with one another and/or with Mlh1-Mlh3 [51].

In this study we created a structure-function map of Mlh3 by analyzing 60 new *mlh3* alleles in *S. cerevisiae*. Five alleles predicted to disrupt the Mlh1-Mlh3 endonuclease motif conferred defects in both MMR and crossing over, providing further support that endonuclease activity is required for both functions. Importantly, we identified five *mlh3* mutations that specifically disrupted MMR, and one mutation that specifically disrupted crossing over. By performing biochemical and genetic analyses of the separation of function Mlh1-mlh3 complexes we suggest that the defects seen in our mutants can be explained by a weakening of protein-protein interactions, which can be tolerated in meiosis, but not MMR. Importantly, our analysis of these mutants revealed unexpected ways in which defective meiotic components can alter the fate of meiotic recombination intermediates.

## Results

### Rationale for site-directed mutagenesis of *MLH3*

Mlh3 contains a highly conserved N-terminal ATP binding motif, a dynamic and unstructured motif known as the linker arm, and an endonuclease active site that overlaps with a C-terminal Mlh1 interaction domain [52]. We performed a clustered charged to alanine scanning mutagenesis of the *S. cerevisiae MLH3* gene to create 60 *mlh3* variants (Fig 2; S1-S3 Tables). Charged residues were considered “clustered” if there were at least two charged residues, consecutive or separated by at most one amino acid, within the primary sequence of Mlh3. Such a directed approach, in the absence of a complete crystal structure, is aimed at targeting the surface of a protein where clusters of charged residues likely reside, while minimizing changes within the interior. In this model, replacement of a charged patch from Mlh3’s surface with alanine residues would disrupt protein-protein or protein-DNA interactions without affecting Mlh3 structure. This unbiased mutagenesis has been successfully applied to study the functional domains of several proteins [53, 54], and has provided a comprehensive view of the functional organization of *MLH1* [55]. As shown below, we identified mutations that caused defects in MMR but not crossing over, likely through disrupted interactions with Mlh1 and other MMR and meiotic CO factors.

**Fig 2.**
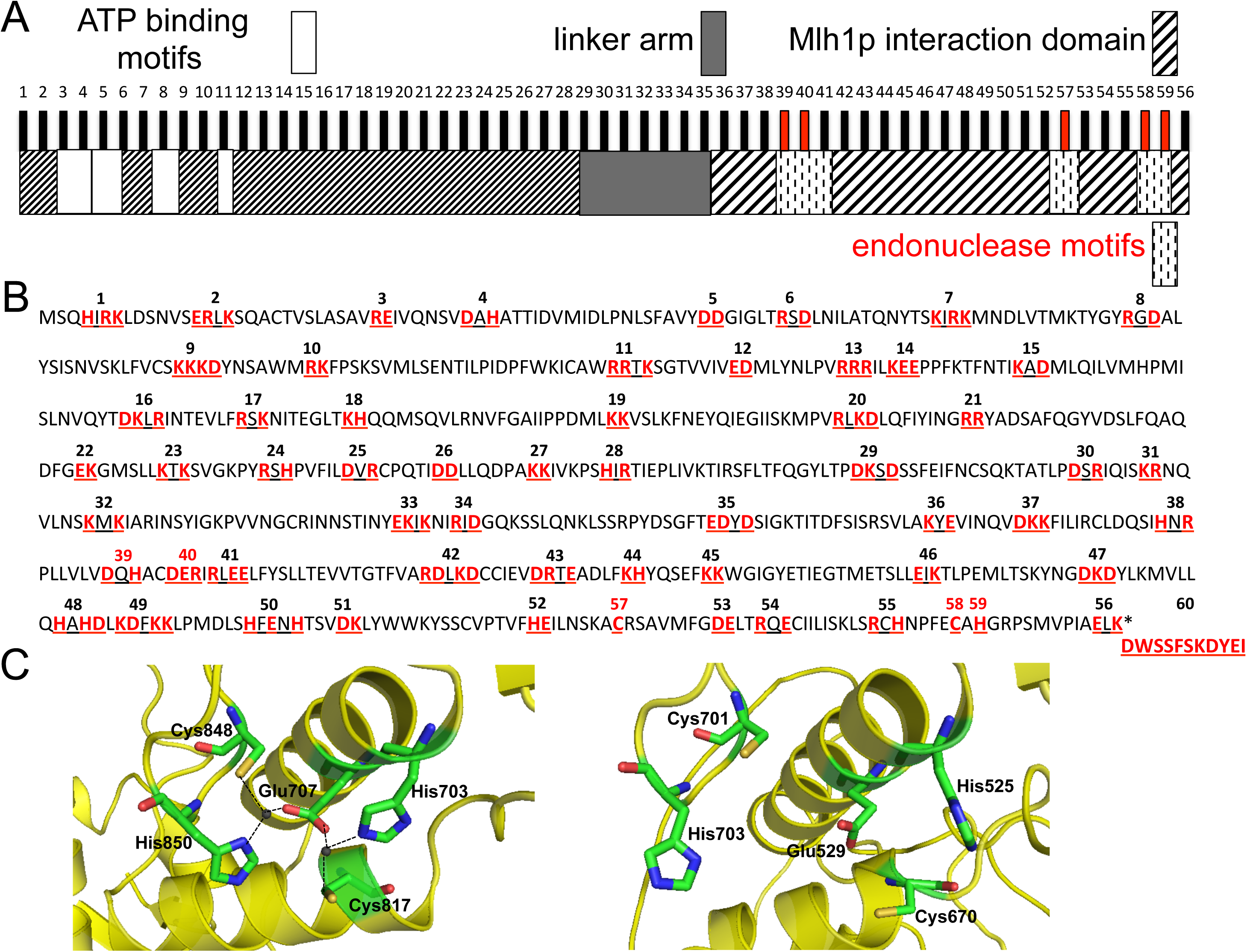
Site directed mutagenesis of *MLH3.* A. Functional organization of Mlh3 based on sequence homology and secondary structure prediction [52]. The vertical bars indicate the approximate position of the *mlh3* mutations (except *mlh3-60*) analyzed in this study and described in panel B. *mlh3-39, −40, −57, −58,* and *-59* colored in red are based on highly conserved residues in the endonuclease motifs of Pms1 which were shown in the crystal structure of Mlh1-Pms1 to form a single metal binding site [52] described in panel C. B. Amino acid positions of charged-to-alanine substitutions presented in red on the primary sequence of *Saccharomyces cerevisiae* Mlh3. Each cluster of underlined residues represents one allele corresponding to the vertical bars in panel A. *mlh3-39, -40, -57, -58,* and *-59* are colored in red as in panel A. *mlh3-60* represents the last 11 residues of Pms1 which constitute patch II of the heterodimerization interface of Mlh1-Pms1 [52]. C. Metal binding site of Pms1 (left panel) from [52] comprised of the five highlighted residues (H703, E707, C817, C848, and H850) were found to be highly conserved in Mlh3 (right panel) based on sequence alignment and structural modeling (H525, E529, C670, C701, and H703) and were targeted in the mutagenesis described in this study (alleles represented in red in A and B).

### Structure-function analysis of Mlh3

We analyzed the effect of *mlh3* mutations on MMR in vegetatively grown cells and on meiotic COs in diploids induced to undergo sporulation. For MMR we employed the *lys2-A_14_* reversion assay to assess the mutation rate in *mlh3* haploid strains (S1 Table; [56]). In this assay the median reversion rate of *mlh3*Δ is six-fold higher than wild-type (Fig 3B; Table 1; [5,6]). To measure meiotic crossing over we crossed mutant *mlh3* strains to *mlh3*Δ strains to form diploids that were then sporulated (S2 Table). The resulting tetrads were directly visualized for chromosome VIII CO events using a spore autonomous fluorescence assay ([57]; Fig 3A). In *mlh3*Δ strains we observed a more than two-fold decrease in crossing over, as measured by percent tetratype, compared to wild-type (Fig 3B). Similar effects of the *mlh3*Δ mutation on crossing over were seen at other genetic intervals [5-8]. It is important to note that nonparental ditype (NPD) events were not scored because they cannot be distinguished from Meiosis I non-disjunction events [57].

**Fig 3.**
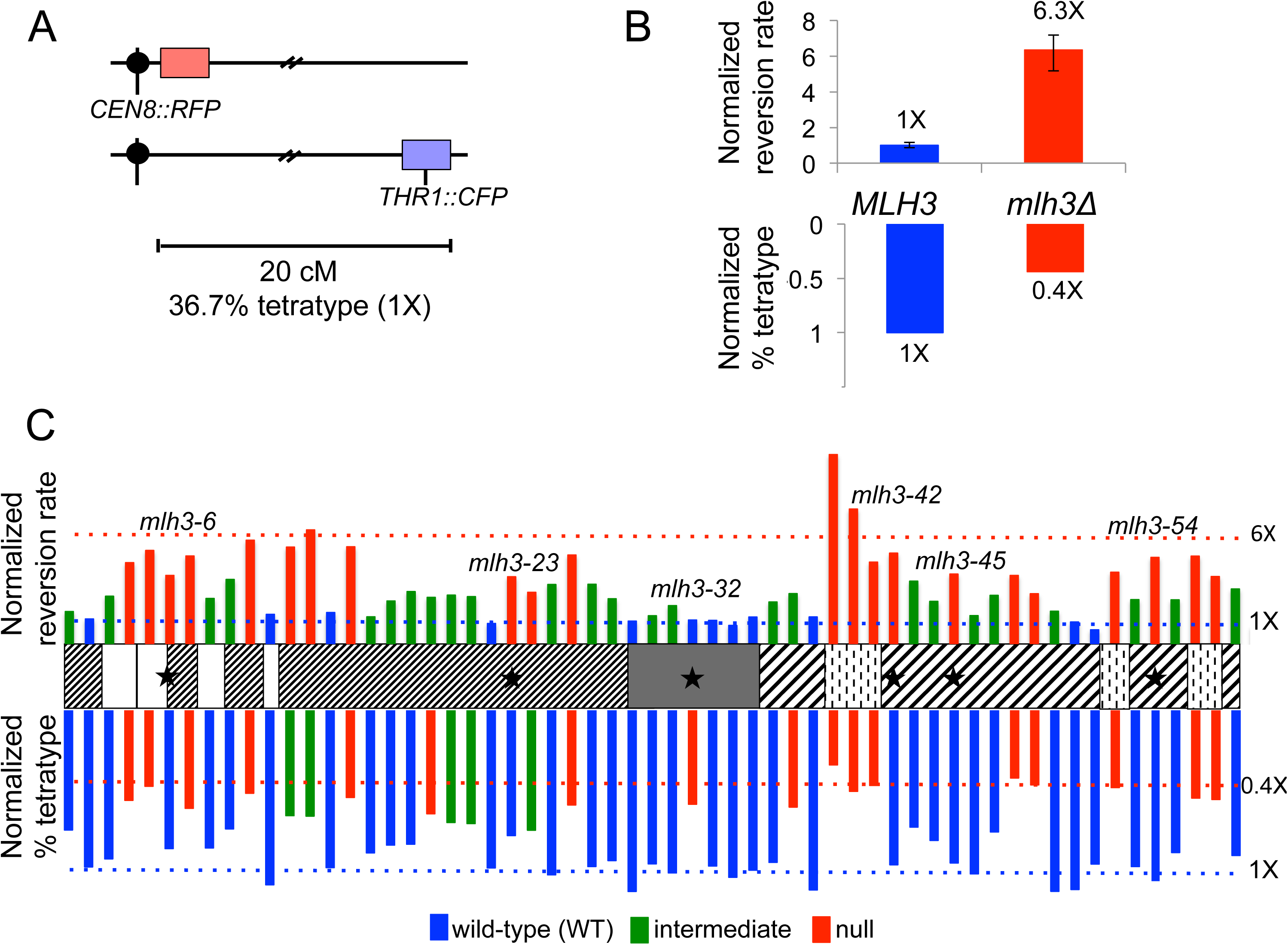
Structure-function map of *Saccharomyces cerevisiae* Mlh3. A. Spore-autonomous fluorescent protein expression was used to quantify crossing over. Shown is the starting parental configuration on chromosome VIII with a map distance of 20 cM separating the red fluorescent protein (RFP) marker and the blue fluorescent protein (CFP) marker [57]. Percent tetratype at this interval in wild-type meiosis is 36.7%. B. Mismatch repair (top) and crossing over (bottom) phenotype of *MLH3* (blue) vs *mlh3Δ* (red). Mismatch repair was measured using the *lys2-A*_*14*_ reversion assay [56] and crossing over was measured using the assay depicted in panel A. Bars represent the median reversion rates (error bars based on 95% confidence intervals) and percent tetratype normalized to *MLH3* (1X). C. The vertical bars indicate the approximate position of the *mlh3* mutations analyzed in this study (except *mlh3-60*) with the height of each bar corresponding to the phenotype relative to *MLH3* (1X). Red indicates a null phenotype, blue indicates wild-type (WT), and green indicates intermediate. For mismatch repair (top), bars represent reversion rates of at least 10 independently tested cultures from two independently constructed strains presented here normalized to *MLH3* median rate 1X=1.43×10^-6^ (n=140). For crossing over (bottom), bars represent percent tetratype of at least 250 tetrads from two independently constructed strains presented here normalized to *MLH3* percent tetratype 1X=36.7% (n= 226). Blue and red dotted lines represent *MLH3* and *mlh3Δ* respectively. ★ indicate separation of function mutants.

**Table 1.** Mismatch repair and crossover phenotypes of the *mlh3* variants as measured in *lys2-A_14_* reversion and spore autonomous fluorescent assays.

| Allele | Reversion rate (x10<sup>-6</sup>) | 95% CI (x10<sup>-6</sup>) | Relative to WT | % tetratype | Relative to WT | Phenotype |  |
| --- | --- | --- | --- | --- | --- | --- | --- |
|  |  |  |  |  |  | MMR | CO |
| *MLH3* | 1.43 | 1.23-1.65 | 1.00 | 36.70 | 1.00 | + | + |
| *mlh3Δ* | 9.07 | 7.4-10.28 | 6.34 | 16.10 | 0.44 | - | - |
| *mlh3-1* | 2.72 | 2.14-3.13 | 1.90 | 27.27 | 0.74 | ± | + |
| *mlh3-2* | 2.09 | 1.63-2.94 | 1.46 | 35.69 | 0.97 | + | + |
| *mlh3-3* | 4.03 | 3.21-5.39 | 2.82 | 33.86 | 0.92 | ± | + |
| *mlh3-4* | 6.89 | 4.16-8.68 | 4.82 | 20.51 | 0.56 | - | - |
| *mlh3-5* | 7.93 | 6.79-9.65 | 5.55 | 17.30 | 0.47 | - | - |
| *mlh3-6* | 5.80 | 2.44-10.7 | 4.06 | 31.52 | 0.86 | - | + |
| *mlh3-7* | 7.46 | 5.59-10.16 | 5.22 | 22.33 | 0.61 | - | - |
| *mlh3-8* | 3.84 | 2.77-5.10 | 2.68 | 31.37 | 0.85 | ± | + |
| *mlh3-9* | 5.46 | 3.78-6 | 3.82 | 27.02 | 0.74 | ± | + |
| *mlh3-10* | 8.80 | 7.72-11.98 | 6.16 | 18.89 | 0.51 | - | - |
| *mlh3-11* | 2.49 | 1.57-2.83 | 1.74 | 39.77 | 1.08 | + | + |
| *mlh3-12* | 8.22 | 6.2-14.44 | 5.75 | 24.00 | 0.65 | - | ± |
| *mlh3-13* | 9.69 | 6.08-25.7 | 6.77 | 24.10 | 0.66 | - | ± |
| *mlh3-14* | 2.64 | 1.57-3.99 | 1.85 | 35.85 | 0.98 | + | + |
| *mlh3-15* | 8.25 | 5.84-9.6 | 5.77 | 19.85 | 0.54 | - | - |
| *mlh3-16* | 2.27 | 1.7-4.78 | 1.59 | 32.49 | 0.89 | ± | + |
| *mlh3-17* | 3.62 | 2.7-6.39 | 2.53 | 30.71 | 0.84 | ± | + |
| *mlh3-18* | 4.42 | 2.74-6.40 | 3.09 | 30.51 | 0.83 | ± | + |
| *mlh3-19* | 3.93 | 3.65-5.23 | 2.75 | 23.57 | 0.64 | ± | - |
| *mlh3-20* | 4.12 | 3.19-6.05 | 2.88 | 25.56 | 0.70 | ± | ± |
| *mlh3-21* | 3.99 | 3.41-5.32 | 2.79 | 25.81 | 0.70 | ± | ± |
| *mlh3-22* | 1.71 | 1.17-2.96 | 1.19 | 35.92 | 0.98 | + | + |
| *mlh3-23* | 5.69 | 4.37-7.56 | 3.98 | 28.54 | 0.78 | - | + |
| *mlh3-24* | 4.37 | 2.57-10.26 | 3.05 | 27.31 | 0.74 | - | ± |
| *mlh3-25* | 5.03 | 4.57-6.22 | 3.52 | 37.50 | 1.02 | ± | + |
| *mlh3-26* | 7.54 | 3.97-12.11 | 5.28 | 21.56 | 0.59 | - | - |
| *mlh3-27* | 5.05 | 2.75-6.78 | 3.53 | 35.60 | 0.97 | ± | + |
| *mlh3-28* | 3.81 | 2.49-4.51 | 2.66 | 34.25 | 0.93 | ± | + |
| *mlh3-29* | 1.91 | 0.61-4.69 | 1.33 | 41.29 | 1.13 | + | + |
| *mlh3-30* | 2.36 | 1.7-3.6 | 1.65 | 35.00 | 0.95 | ± | + |
| *mlh3-31* | 3.23 | 2.01-6.91 | 2.26 | 37.06 | 1.01 | ± | + |
| *mlh3-32* | 2.00 | 1.54-2.22 | 1.40 | 21.40 | 0.58 | + | - |
| *mlh3-33* | 1.97 | 1.65-2.58 | 1.38 | 35.49 | 0.97 | + | + |
| *mlh3-34* | 1.54 | 1.25-2.82 | 1.07 | 38.06 | 1.04 | + | + |
| *mlh3-35* | 2.26 | 0.65-3.21 | 1.58 | 36.43 | 0.99 | + | + |
| *mlh3-36* | 3.54 | 2.37-5.45 | 2.47 | 34.66 | 0.94 | ± | + |
| *mlh3-37* | 4.25 | 2.54-5.15 | 2.97 | 22.04 | 0.60 | ± | - |
| *mlh3-38* | 2.26 | 1.14-3.94 | 1.58 | 40.94 | 1.12 | + | + |
| *mlh3-39* | 16.11 | 10.54-19 | 11.26 | 12.42 | 0.34 | - | - |
| *mlh3-40* | 11.46 | 5.81-16.57 | 8.02 | 18.45 | 0.50 | - | - |
| *mlh3-41* | 6.94 | 3.56-9.1 | 4.85 | 17.11 | 0.47 | - | - |
| *mlh3-42* | 7.70 | 5.38-12 | 5.39 | 35.20 | 0.96 | - | + |
| *mlh3-43* | 5.31 | 4.09-7.23 | 3.71 | 26.61 | 0.73 | ± | + |
| *mlh3-44* | 3.59 | 2.87-4.25 | 2.51 | 29.65 | 0.81 | ± | + |
| *mlh3-45* | 5.92 | 3.49-11.7 | 4.14 | 34.79 | 0.95 | - | + |
| *mlh3-46* | 2.38 | 1.75-3.01 | 1.66 | 37.27 | 1.02 | ± | + |
| *mlh3-47* | 4.10 | 2.98-5.29 | 2.87 | 27.72 | 0.76 | ± | + |
| *mlh3-48* | 5.80 | 4.01-8.93 | 4.06 | 15.41 | 0.42 | - | - |
| *mlh3-49* | 4.24 | 3.6-9.72 | 2.97 | 16.98 | 0.46 | - | - |
| *mlh3-50* | 2.75 | 2.24-3.33 | 1.92 | 41.26 | 1.12 | ± | + |
| *mlh3-51* | 1.83 | 1.04-3.23 | 1.28 | 40.84 | 1.11 | + | + |
| *mlh3-52* | 1.17 | 0.79-2.7 | 0.82 | 35.04 | 0.95 | + | + |
| *mlh3-57* | 6.07 | 4.74-9.4 | 4.25 | 17.60 | 0.48 | - | - |
| *mlh3-53* | 3.74 | 2.52-6.9 | 2.62 | 35.61 | 0.97 | ± | + |
| *mlh3-54* | 7.34 | 5.59-9.97 | 5.13 | 38.93 | 1.06 | - | + |
| *mlh3-55* | 3.72 | 2.22-5.45 | 2.60 | 32.41 | 0.88 | ± | + |
| *mlh3-58* | 7.45 | 5.23-10.63 | 5.21 | 20.00 | 0.54 | - | - |
| *mlh3-59* | 5.71 | 4.28-8.19 | 3.99 | 20.31 | 0.55 | - | - |
| *mlh3-56* | 4.65 | 3.53-6.11 | 3.25 | 33.09 | 0.90 | ± | + |
| *mlh3-60* | 2.50 | 1.18-4.45 | 1.75 | 33.83 | 0.92 | + | + |

Two independently constructed strains with *mlh3* variants were analyzed in the EAY3255 background which contains the *lys2::insE-A*_*14*_ for MMR testing and the red fluorescent protein for meiotic testing. Haploid strains were examined for reversion to Lys^+^. At least n=10 reversion assays were performed per allele. Median reversion rates are presented with 95% confidence intervals (CI), and relative reversion rates compared with the wild-type strain are shown. The haploid strains were mated to EAY3486, which contains the blue fluorescent protein to make diploids suitable for meiotic testing. Diploid strains were induced for meiosis and % tetratype was measured. At least n=250 tetrads were counted for each allele. WT, wild-type. +, indistinguishable from WT as measured by 95% CI (for reversion rates) or χ2 (p<0.01, for % tetratype). -, indistinguishable from null as measured by 95% CI or χ2 (p<0.01). +/-, distinguishable from both wild-type and null as measured by 95% CI or χ2 (p<0.01).

Similar to work performed on a smaller number of *mlh3* alleles and a structure-function analysis of *MLH1*, we found that *MLH3* MMR functions were more sensitive to mutagenesis than CO functions (Fig 3C; [6, 55]). Phenotypes exhibited by *mlh3* strains containing mutations in the ATP-binding motif suggested that this region plays a more critical role in MMR compared to crossing over. However a region just beyond the ATP-binding domain appeared insensitive to mutagenesis. A null phenotype for both functions was observed in strains bearing mutations in endonuclease motifs, further confirming that endonuclease activity is essential for MMR and crossing over (Fig 3C; Table 1; [5, 8, 16]).

We showed that the two functions of Mlh3 are genetically separable. Comparison of the MMR and CO assay results for each individual allele led to the identification of six separation of function mutations in the ATP-binding motifs, the N-terminal domain beyond the ATP-binding motifs, the linker arm, and the interaction domain (Fig 3C, indicated by stars). One of these alleles (*mlh3-32*) conferred a nearly wild-type phenotype for MMR and a null phenotype for crossing over on chromosome VIII (hereafter referred to as MMR^+^, CO^-^). The remaining five separation of function mutations (*mlh3-6*, *mlh3-23*, *mlh3-42*, *mlh3-45*, and *mlh3-54*) conferred null MMR phenotypes and nearly wild-type levels of crossing over (hereafter referred to as MMR^-^, CO^+^). The null phenotype observed in the separation of function mutants may result from a defect in DNA binding/substrate specificity, endonuclease activity, interactions with specific MMR and meiotic CO factors, or changes in protein conformation. It is important to note that a co-crystal structure of the N-terminal domain of *E. coli* MutL (LN40) and *E. coli* MutS was recently solved [58]. This work showed that conformational changes license MutS-MutL interaction and are essential for MMR.

### The endonuclease active sites in Mlh3 and Pms1 appear to be similar

*S. cerevisiae* Mlh1-Pms1 and Mlh1-Mlh3 and human MLH1-PMS2 display latent endonuclease activities dependent on the integrity of a highly conserved metal binding motif DQHA(X)_2_E(X)_4_E [14-17]. This motif is critical for Mlh3’s MMR and meiotic functions [5]. Two additional motifs were implicated in MLH family endonuclease function based on sequence alignment: ACR and C(P/N)HGRP [59]. In the Mlh1-Pms1 C-terminal domains crystal structure, five Pms1 residues, located in the three conserved motifs (H703, E707, C817, C848, H850), form a metal binding site through folding of the Pms1 C-terminal domain (Fig 2C; [52]). This organization was also seen in the crystal structure of the C-terminal domain of *B. subtilis* endonuclease MutL (all but H703 are conserved; [60]).

We performed a sequence alignment of Mlh3, Pms1, and *B. subtilis* MutL and found all three possess conserved metal binding motifs, with the following five residues predicted to form the endonuclease active site in Mlh3: H525, E529, C670, C701, and H703. In addition, we constructed a homology model of *S. cerevisiae* Mlh3, and found the C-terminal domain can potentially fold in a similar manner to Pms1 such that these five conserved residues form a single putative metal binding site (Fig 2C). These residues were targeted for site directed mutagenesis of *MLH3* (Fig 2B, numbered in red). As shown in Fig 3C (represented in the C-terminal domain by dotted white squares) and Fig 4A and Table 1, mutations in the putative conserved metal binding motifs of Mlh3 (*mlh3-39(D523A, H525A), −40(D528A, E529A, R530A), −57(C670A), −58(C701A), and -59(H703A)*) conferred null phenotypes for MMR and crossing over, indicating they are essential for Mlh3 function. Thus these genetic data, combined with the high sequence homology, suggest that H525, E529, C670, C701, and H703 in the C-terminal domain of Mlh3 form the catalytic active site (Fig 2C).

**Fig 4.**
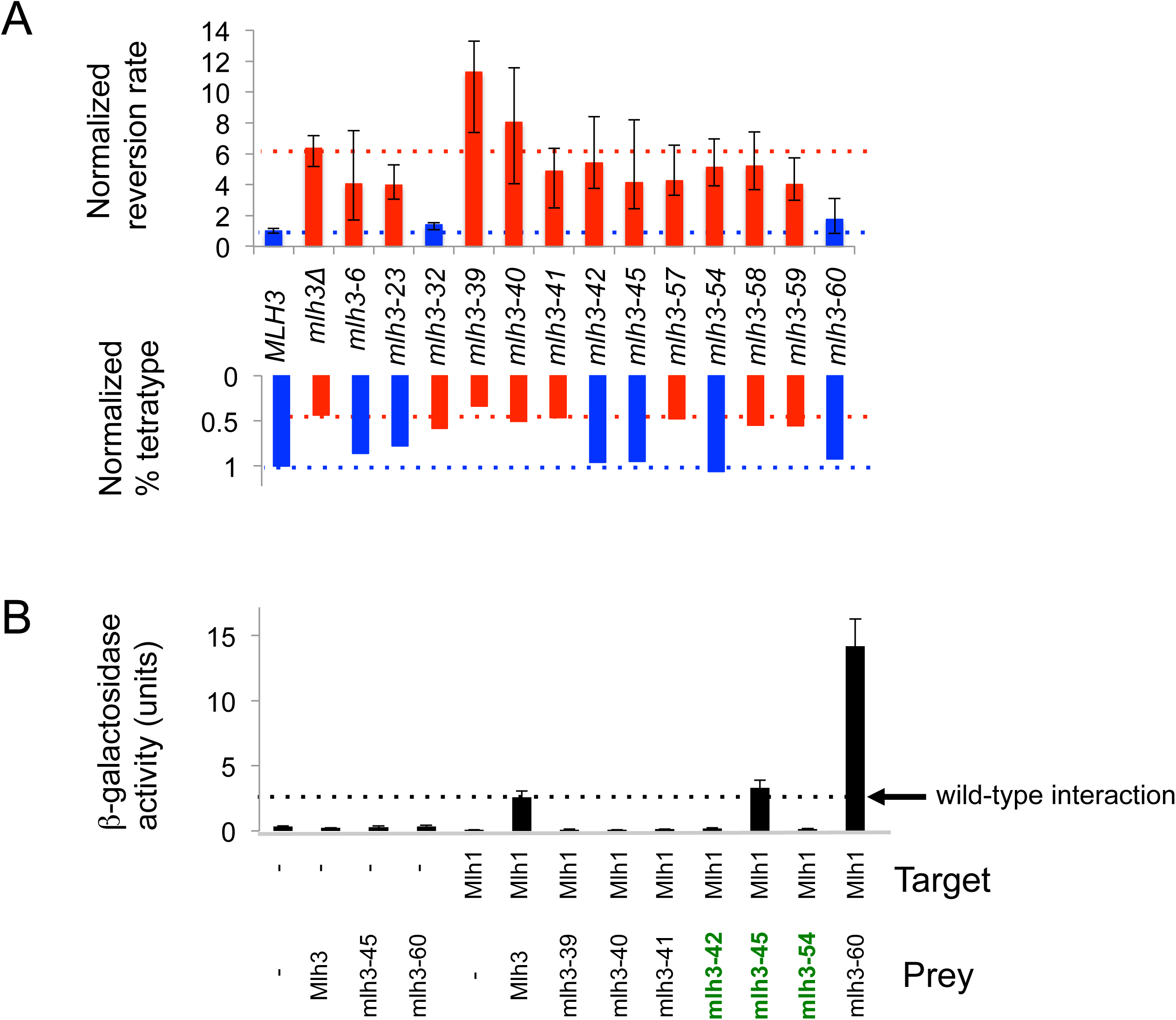
*mlh3-42, −54* weaken Mlh1 interaction yet maintain crossover function. A. MMR and CO phenotypes for separation of function, endonuclease, and C-terminal tail mutants (*mlh3-60*). B. Yeast two-hybrid interactions between lexA-Mlh1 (target) and Gal4-Mlh3 (amino acids 481-715; prey) or Gal4-mlh3-39, −40, −41, −42, −45, −54, −60 derivative constructs, as measured in the ONPG assay for β-galactosidase activity. Error bars indicate standard error of mean from at least three independent assays. *mlh3* separation of function alleles indicated in green font.

The endonuclease motifs in Mlh3 overlap with the C-terminal Mlh1 interaction domain. To determine if mutations in the Mlh3 endonuclease motifs disrupted interaction with Mlh1, three alleles spanning the DQHA(X)_2_E(X)_4_E endonuclease motif (*mlh3-39, −40, and −41*) were analyzed by yeast two-hybrid for interaction with Mlh1. We also tested these alleles because a previously characterized mutation in the DQHA(X)_2_E(X)_4_E endonuclease motif (*mlh3-E529K*) disrupts Mlh1-Mlh3 interactions [5]. As shown in Fig 4B, these mutations disrupted Mlh1-Mlh3 interactions, possibly by altering the endonuclease active site structure. This idea is supported by the Mlh1-Pms1 crystal structure. In this model, heterodimer stability is maintained through interactions between the C-terminal domain of Mlh1 and the endonuclease active site of Pms1 [52]. We cannot rule out the possibility that the null phenotypes observed for MMR and crossing over in *mlh3-39, −40, and −41* were caused by specifically mutating residues that comprise the Mlh1-Mlh3 dimerization interface without causing a gross disruption in protein folding.

The Mlh1-Pms1 C-terminal domain structure reveals three patches constituting the heterodimerization interface of Mlh1-Pms1 [52]. Patch I is a pseudosymmetric hydrophobic core, Patch II is composed of the last 12 residues of Pms1 and contributes two salt bridges, and Patch III involves the C-terminus of Mlh1 and contributes to the Pms1 metal binding site [52]. Patches I and III are likely maintained in the Mlh1-Mlh3 heterodimerization interface, but Mlh3 lacks the last 11 residues that comprise the bulk of Patch II. This finding gives a likely explanation for partial disruption of the Mlh1-Mlh3 complex when we attempted to analyze it further by gel-filtration [16]. We hypothesized that restoring Patch II to the Mlh1-Mlh3 interaction interface will strengthen this interaction. We engineered a fusion construct of Mlh3 carrying the last 11 residues of Pms1 (*mlh3-60*, Fig 2B in red). As shown in Fig 4B, when we inserted the last 11 residues of Pms1 after the C-terminal residue of Mlh3, we observed a striking increase in the strength of the interaction between Mlh1 and Mlh3 as measured in the yeast two-hybrid assay (2.6 ± 0.5 Miller units of β-galactosidase activity for wild-type Mlh1-Mlh3 compared to 14.2 ± 2.1 for Mlh1-mlh3-60). We initially hypothesized that such an enhanced interaction would be detrimental to MMR because it would sequester Mlh1 from the major MMR endonuclease Pms1. Surprisingly we did not observe a significant effect of the *mlh3-60* mutation on MMR or on crossing over (Fig 4A), suggesting that strengthening the interaction between Mlh1-Mlh3 does not affect formation of the Mlh1-Pms1 heterodimer.

### *mlh3-42* and *mlh3-54* weaken interaction with Mlh1 yet maintain wild-type levels of crossing over

Three of the five MMR^-^, CO^+^ alleles (*mlh3-42, −45*, and *-54*) contained mutations that mapped to the Mlh1 interaction interface. We performed a two-hybrid assay to test whether these mutations affected Mlh1-Mlh3 dimerization. The *mlh3-45* mutation did not alter Mlh1-Mlh3 interactions; however both the *mlh3-42* and *mlh3-54* mutations disrupted this interaction (Fig 4B). While such a result could explain the null MMR phenotype conferred by *mlh3-42* and *mlh3-54*, it does not explain why these strains are functional for meiotic crossing over. One explanation is that additional pro-CO factors act as structural scaffolds to stabilize the weakened Mlh1-mlh3 heterodimer, thus allowing it to perform its function at dHJs. Several observations support this idea: 1. The pro-CO factors Msh4-Msh5, Sgs1-Top3-Rmi1, Zip3, and Exo1 interact with one another and/or with Mlh1-Mlh3 [29, 40, 41, 42, 61, 62]. 2. Studies in mice showed that MLH1 and MLH3 do not form a complex until mid to late pachytene; at early to mid pachytene, only MLH3 foci are seen [37, 44]. 3. Exo1’s role in crossing over is independent of its enzymatic activity; it is suggested to play a structural role, acting as a platform for pro-CO factors [29]. Together these observations support the presence of a resolvase complex at CO sites that regulates the endonuclease activity of Mlh1-Mlh3 (see Discussion). Alternatively, a weak Mlh1-Mlh3 interaction defect is sufficient to inhibit a yeast-two hybrid interaction, but not affect meiotic recombination if the strength of the Mlh1-Mlh3 interaction is not a limiting factor for CO resolution.

### Mlh1-mlh3-32 and Mlh1-mlh3-45 display wild-type endonuclease activities but only Mlh1-mlh3-32 endonuclease is stimulated by Msh2-Msh3

We examined Mlh1-mlh3 mutant complexes for endonuclease activity [16, 17], focusing on opposite separation of function mutants Mlh1-mlh3-32 (MMR^+^, CO^-^) and Mlh1-mlh3-6 and Mlh1-mlh3-45 (MMR^-^, CO^+^). Mlh1-mlh3-45, located in the C-terminal Mlh1 interaction domain, was chosen because it is the only separation of function mutant in that domain that displayed wild-type Mlh1-Mlh3 interactions as measured in the two-hybrid assay (Fig 4B). As shown in Fig 5 and S1 Fig, all three mutant complexes purified as heterodimers and display endonuclease activities similar to wild-type. When we initiated this work we thought that separation of function mutant complexes might show endonuclease defects indicating that this activity is more critical for MMR or crossing over, or show no endonuclease defects because mutant complexes were defective in interacting with MMR or CO specific factors. Our finding that all three mutants have enzymatic activity comparable to wild-type is consistent with the interaction defect model (see below). The *mlh3-6* mutation maps close to conserved sites in the ATP binding motif (Fig 3C). For this reason we tested whether the mutant complex displayed a defect in ATPase activity. As shown in S1C Fig, Mlh1-Mlh3 and Mlh1-mlh3-6 displayed similar ATPase activities.

**Fig 5.**
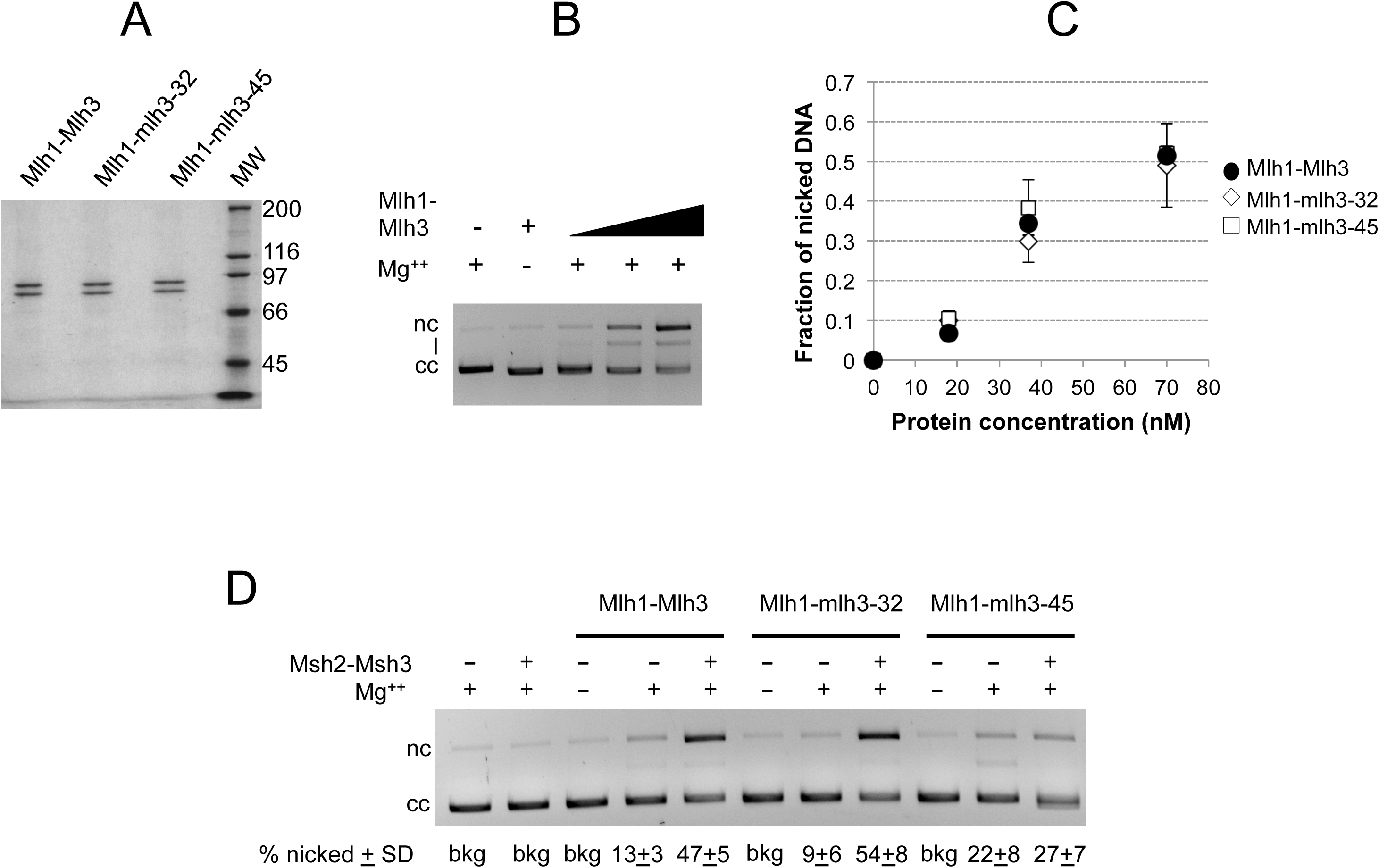
Mlh1-mlh3-32 and Mlh1-mlh3-45 display wild-type endonuclease activities that are differentially stimulated by Msh2-Msh3. A. SDS-PAGE analysis of purified Mlh1-Mlh3, Mlh1-mlh3-32 and Mlh1-mlh3-45. Coomassie Blue R250-stained 8% Tris-glycine gel. 0.5 μg of each protein is shown. MW = Molecular Weight Standards from top to bottom- 200, 116, 97, 66, 45 kD). B, C. Mlh1-Mlh3, Mlh1-mlh3-32 and Mlh1-mlh3-45 (18, 37, 70nM) were incubated with 2.2 nM supercoiled pBR322 DNA, and analyzed in agarose gel electrophoresis (C) and the endonuclease activity was quantified (average of 6 independent experiments presented +/-SD) as described in the Experimental Procedures. Ladder: 1 kb DNA ladder (New England Biolabs). Migration of closed circular (cc), nicked (nc) and linear (l) pBR322 DNA is indicated. D. Endonuclease assays were performed as in B., but contained 20 nM of the indicated wild-type or mutant Mlh1-Mlh3 complex and 40 nM Msh2-Msh3 when indicated. Reactions were performed in triplicate, samples were resolved on agarose gels, and the fraction of nicked DNA was quantified, averaged, and the standard deviation between experiments was calculated. The average fraction of supercoiled substrate cleaved is presented +/-S.D. below the gel. (bkg) background, (cc) closed circular DNA, (nc) nicked DNA.

Because Mlh1-Mlh3’s endonuclease activity is enhanced by Msh2-Msh3 [16], we tested whether the opposite separation of function phenotypes of Mlh1-mlh3-32 and Mlh1-mlh3-45 could be explained by defective interactions with MSH complexes. As shown in Fig 5D, Mlh1-mlh3-32 endonuclease activity but not Mlh1-mlh3-45 could be stimulated by Msh2-Msh3. These data are consistent with the MMR^-^, CO^+^ phenotype exhibited by *mlh3-45* mutants resulting from a defect in interacting with the MMR component Msh2-Msh3. However, the MSH-MLH interaction is likely maintained in *mlh3-32*, suggesting that this mutant may be defective in interactions with other meiotic factors. In a first step towards testing if *mlh3-32* mutants are defective in interactions with meiotic CO factors, we determined if the *mlh3-32* mutation is dominant. Such a phenotype could provide additional hints on the nature of the *mlh3-32* meiotic defect. We mated EAY3552 (*mlh3-32, CEN8Tomato::LEU2*) to EAY3339 (relevant genotype *MLH3, THR1::m-Cerulean-TRP1*), and the sporulated progeny displayed a tetratype frequency similar to wild-type (40.9%, n = 252 tetrads), indicating that *mlh3-32* is recessive.

### Cumulative genetic distance and spore viability measurements confirm fluorescent assay results for *mlh3* separation of function alleles

The meiotic CO phenotype for the separation of function mutants was determined at four consecutive intervals in chromosome (XV) using traditional genetic map distance analyses (Fig 6A; [5-7]). The overall effect of *mlh3* mutations in crossing over on chromosome XV was similar to that determined on chromosome VIII (Fig 3C, top panel, indicated with stars; Fig 6B; S4 Table). *mlh3-6*, *mlh3-42*, *mlh3-45*, and *mlh3-54* appear similar to wild-type; *mlh3-23* displays an intermediate phenotype, and *mlh3-32* appears similar to the null (Fig 6B; S4 Table). Thus, these mutants were confirmed as separation of function alleles and are candidates for in-depth characterization and high-resolution recombination mapping.

**Fig 6.**
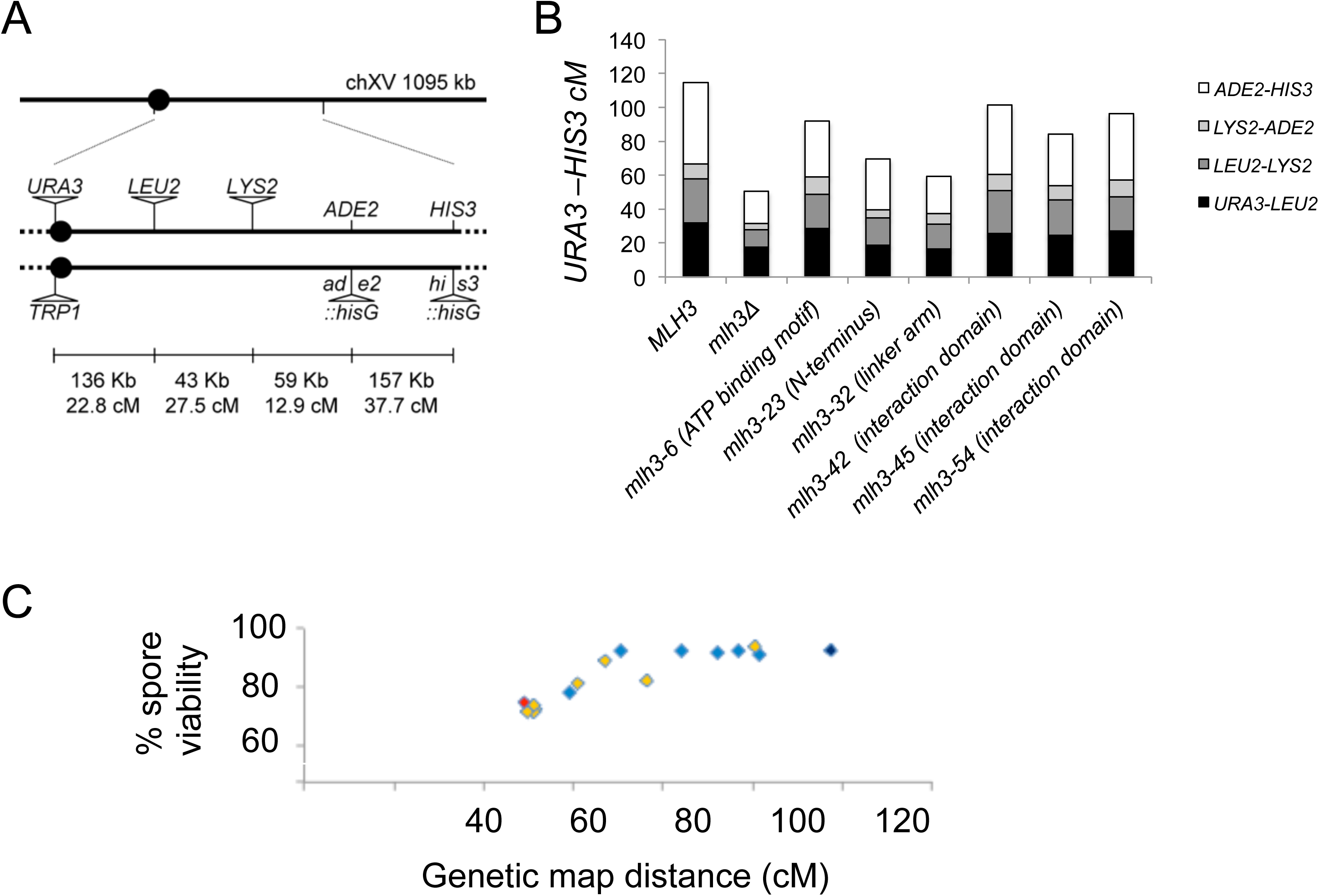
Cumulative genetic distance and spore viability of *mlh3* separation of function mutants. A. Distribution of genetic markers on chromosome XV used to determine genetic distances in the EAY1112/EAY2413 background (S1 Table). The solid circle indicates the centromere. The distances between markers are not drawn to scale. The actual physical and genetic distances in the wild-type diploid are given numerically for each interval and for the entire region between *CENXV* and *HIS3* [7]. B. Cumulative genetic distances between *URA3* and *HIS3* markers from tetrads of *MLH3* and indicated *mlh3* variants. Each bar is further divided into sectors that correspond to the four genetic intervals that span *URA3-HIS3*. C. Spore viabilities are plotted vs. genetic map distances from panel B for *MLH3* (dark blue), *mlh3Δ* (red), and the separation of function mutants (light blue). Yellow diamonds represent data from Sonntag Brown et al. [6].

It is important to note that the spore viability and genetic distance measurements of the six separation of function alleles indicated that CO levels, represented by genetic distance measurements in Chr. XV, can be reduced to ~69 cM without compromising spore viability (Fig 6C; Table 2). A similar observation was made with *msh4/5* hypomorph alleles where crossing over in the four same intervals in Chr. XV could be reduced to ~50 cM without affecting spore viability [63-64]. Together, these observations and the high resolution mapping below indicate that the baker's yeast meiotic cell does not require the full amount of COs maintained by CO homeostasis (~90; see [65]) for accurate chromosome segregation and to form viable spores. Data were obtained from Table 1 (% tetratype, reversion rate), S4 Table (% spore viability (SV), genetic distance in cM), and Fig 4B (β-gal units in the two-hybrid assay). ND, not determined.

**Table 2.** Summary of *mlh3* separation of function phenotypes.

| Genotype | %SV | Genetic map |  | Reversion rate | β-gal units |
| --- | --- | --- | --- | --- | --- |
|  |  | distance (cM) | % tetratype | X10<sup>-6</sup> (95%CI) | (±SEM) |
| *MLH3* | 93.3 | 114.5 | 36.7 | 1.4 (1.2-1.7) | 2.6 (±0.5) |
| *mlh3Δ* | 74.9 | 50.5 | 16.1 | 9.1 (7.4-10.3) | ND |
| *mlh3-6* | 91.4 | 91.9 | 31.5 | 5.8 (2.4-10.7) | ND |
| *mlh3-23* | 90.9 | 69.6 | 28.5 | 5.7 (4.4-7.6) | ND |
| *mlh3-32* | 78.5 | 59.2 | 21.4 | 2.0 (1.5-2.2) | ND |
| *mlh3-42* | 91.8 | 101.3 | 35.2 | 7.7 (5.4-12.0) | 0.20 (±0.04) |
| *mlh3-45* | 92.9 | 84.2 | 34.8 | 5.9 (3.5-11.7) | 3.3 (±0.6) |
| *mlh3-54* | 92.3 | 96.4 | 38.93 | 7.34 (5.59-9.97) | 0.16 (±0.03) |

### Sgs1 overexpression differentially affects spore viability in mlh3Δ vs. MLH3 and mlh3-D523N strains

As presented in Fig 1, the STR complex can act as both a negative and positive regulator of CO formation in meiotic prophase [8, 30, 31, 46, 50]. In its role as a negative regulator, STR is thought to prevent the formation of aberrant recombination structures by disassembling branched recombination intermediates to form early NCOs via synthesis dependent strand annealing (SDSA), or by re-forming the DSB intermediate. In its role as a pro-CO factor STR promotes stabilization of ZMM complexes on recombination intermediates, leading to the resolution of dHJs by an interference-dependent CO pathway (class I) that requires the Mlh1-Mlh3 endonuclease. In *sgs1Δ* mutants COs have been shown to be ZMM independent [66]. Strand invasion intermediates that escape STR disassembly are thought to be resolved as COs or NCOs using an alternative interference-independent CO pathway (class II) that involves the structure-specific nucleases (SSNs), Mus81-Mms4 and Yen1.

To test for genetic interactions between *SGS1* and *MLH3*, we expressed *SGS1* via its native promoter on a *2μ* multi-copy vector. Sgs1 overexpression enhanced the *mlh3*Δ spore viability defect (S2 Fig: 76% in *mlh3Δ+2μ* vs. 57% in *mlh3Δ+SGS1-2μ*) and conferred a more extreme MI nondisjunction pattern (an excess of 4, 2, 0 viable spore tetrads compared with 3 and 1 viable tetrads ([6]; S2 Fig). However, no such effect was seen in *MLH3* (96% in *MLH3+2μ* vs. 95% in *MLH3+SGS1-2μ*), *mlh3-32* (76% in *mlh3-32+2μ* vs. 84% in *MLH3+SGS1-2μ*) and *mlh3-D523N* strains (S2 Fig: 82% in *mlh3-D523N+2μ* vs. 79% in *mlh3-D523N+SGS1-2μ*). The previously characterized *mlh3-D523N* mutation contains an aspartic acid to asparagine substitution in the DQHA(X)_2_E(X)_4_E metal binding motif of Mlh3. This mutation does not disrupt formation of the Mlh1-Mlh3 complex; however, it conferred a null phenotype for *MLH3* functions in MMR and meiotic CO assays, and the Mlh1-mlh3-D523N complex is defective for endonuclease activity [5, 16]. Nonetheless, this mutant did not behave like *mlh3*Δ in response to Sgs1 overexpression. This finding encouraged us to explore roles for Mlh1-Mlh3 that appear independent of its enzymatic activity.

### High-resolution recombination maps illustrate unexpected effects of *mlh3* hypomorphs and the *mlh3-D523N* allele on resolving meiotic recombination intermediates

We characterized two alleles with opposite separation of function phenotypes, *mlh3-23* (MMR^-^, CO^+^) and *mlh3-32* (MMR^+^, CO^-^), by mapping recombination events genome-wide using the S288c/YJM789 hybrid [67]. We also analyzed the *mlh3-D523N* mutation described above [5, 16]. The Mlh1 protein sequence has two amino acid differences between SK1 and YJM789 strains and three amino acid differences between SK1 and S288c strains. The SK1 Mlh3 protein has 11 amino acid differences with respect to S288c Mlh3 and seven with respect to YJM789 Mlh3. Therefore we analyzed the SK1 *mlh3* mutations in the presence of SK1 *MLH1* in the S288c/YJM789 hybrid to avoid genetic incompatibilities between Mlh1 and Mlh3.

The SK1 *MLH3*, *mlh3-23, mlh3-32*, and *mlh3-D523N* alleles were introduced into an *mlh1Δ* S288c strain and the SK1 *MLH1* allele was introduced into an *mlh3Δ* YJM789 strain (Fig 7A, S1 Table). The SK1 *MLH1, MLH3*, and *mlh3-23*, *mlh3-32*, and *mlh3-D523N* alleles were analyzed as heterozygotes over their respective null mutations (S2 Table; [64]). The spore viabilities of the mutants, *mlh3-23* (84%), *mlh3-32* (82%), and *mlh3-D523N* (82%) were similar to *mlh3Δ* (85%) and the wild-type hybrid (84%; Table 3). Why do wild-type and *mlh3* strains show similar viability in the S288c/YJM789 hybrid? *mlh3Δ* mutants display a range of spore viabilities (70 to 92%) that appear to depend on strain background [5, 6, 68]. This is likely to be a partial explanation; however another study suggested that sequence divergence present in the strain hybrids can affect spore viability through mismatch repair mechanisms that act on heteroduplex DNA formed during genetic recombination [69].

**Fig 7.**
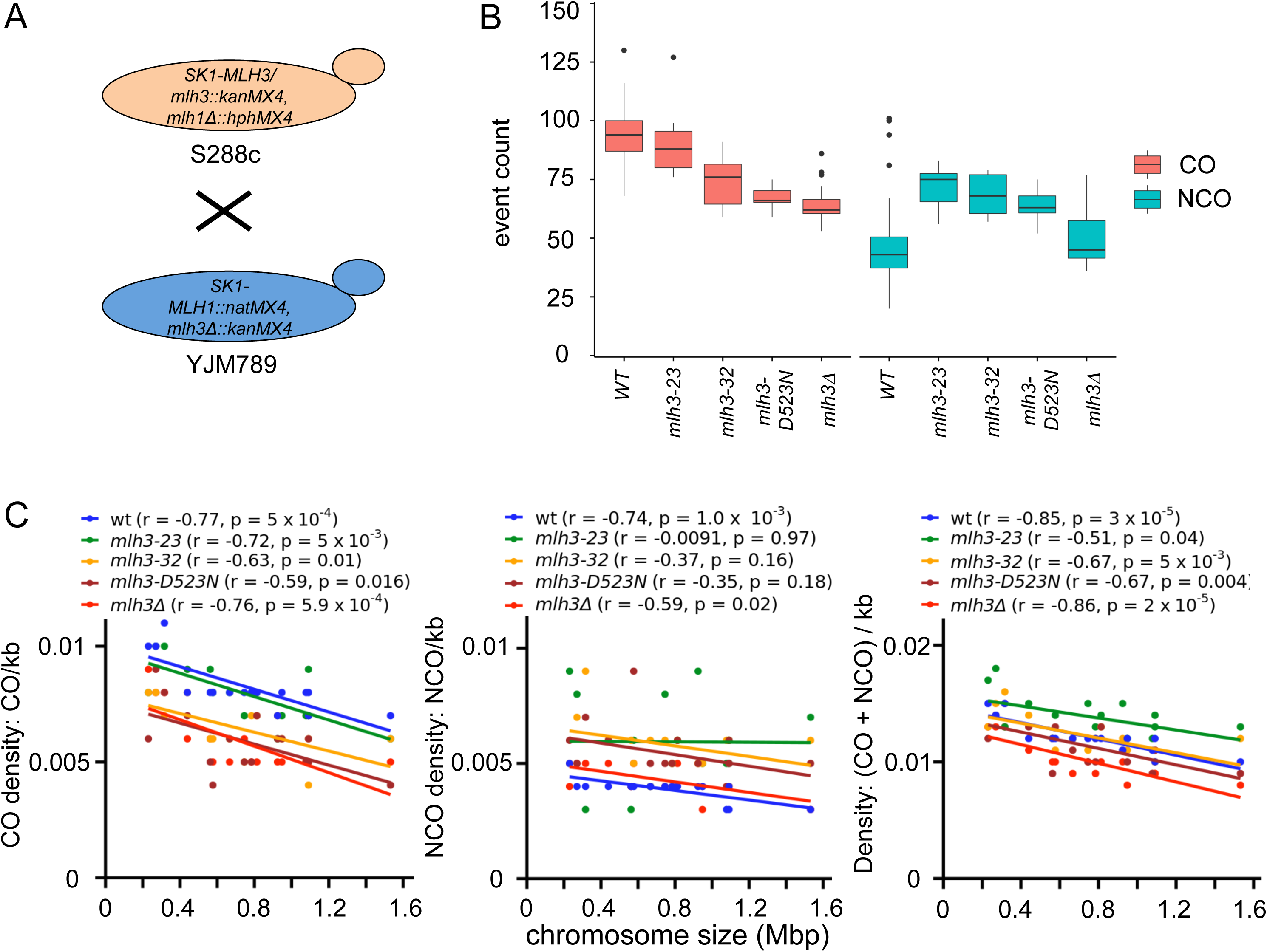
Genome-wide increase in non-crossovers in *mlh3-23, mlh3-32* and *mlh3-D523N* mutants. A. Generation of S288c/YJM789 strains with SK1 *MLH1, MLH3* and the *mlh3-23, mlh3-32, mlh3-D523N* mutant alleles. The SK1 *MLH1, MLH3, mlh3-23, mlh3-32* and *mlh3-D523N* constructs were introduced into *S. cerevisiae* S288c or YJM789 strains by homologous recombination. B. Crossover (CO) and non-crossover (NCO) counts per meiosis for wild-type, *mlh3-23, mlh3-32, mlh3-D523N*, and *mlh3Δ*. The minimum, first quantile, median, third quantile and maximum count are indicated in the box plot. C. Density plot for crossovers (left), non-crossovers (middle) and crossovers plus non-crossovers (right) as a function of chromosome size in wild-type (wt), *mlh3-23, mlh3-32, mlh3-D523N* and *mlh3Δ*.

**Table 3.** Spore viability, crossover (CO) and non-crossover (NCO) values for *mlh3-23*, *mlh3-32, mlh3-D523N*, and *mlh3Δ* mutants in the S288c/YJM789 hybrid.

| Genotype | N | % S.V | Tetrads genotyped | Avg. CO counts ± S.D (Median) | Avg. NCO counts ± S.D (Median) |
| --- | --- | --- | --- | --- | --- |
| S288c x YJM789 | 180 | 84 | 66* | 93 ± 11 (94) | 46 ± 16 (43) |
| S288c x YJM789 with SK1- *MLH1/MLH3* | 88 | 80 | 4 | 95 ± 16<br>(91) | 49 ± 9<br>(48) |
| S288c x YJM789 *mlh3-23* | 80 | 84 | 7 | 92 ± 18<br>(88) | 71 ± 9<br>(75) |
| S288c x YJM789 *mlh3-32* | 80 | 82 | 7 | 74 ± 12<br>(76) | 68 ± 9<br>(68) |
| S288c x YJM789 *mlh3-D523N* | 88 | 82 | 10 | 67 ± 5<br>(66) | 64 ± 7<br>(63) |
| S288c x YJM789 *mlh3Δ* | 120 | 85 | 19<sup>#</sup> | 64 ± 9<br>(62) | 50 ± 12<br>(45) |

N = number of tetrads dissected for measuring spore viability (S.V), S.D = standard deviation.
\*Merged data set from [64, 67]. <sup>#</sup>Data set from Chakraborty et al. [70].

Seven four-spore viable tetrads of *mlh3-23* and *mlh3-32* mutants were sequenced along with ten tetrads of *mlh3-D523N* and four tetrads of the S288c/YJM789 hybrid bearing SK1-*MLH1/MLH3* alleles (S5 Table). The sequence data are available from the National Centre for Biotechnology Information Sequence Read Archive under the accession number SRP096621. The recombination parameters for the wild-type hybrid were generated from a merged set of 66 tetrads from previously published studies [70]. The S288c/YJM789 hybrid bearing SK1-*MLH1/MLH3* alleles showed CO and NCO counts similar to wild-type, indicating that the SK1-*MLH1/MLH3* alleles are functional in the S288c/YJM789 hybrid (Table 3). The segregation of the SNPs in all 28 tetrads is shown in S1 File. The CO and NCO counts for all tetrads and the average CO and NCO counts per chromosome are shown in S6 and S7 Tables, respectively.

As described below, high-resolution recombination mapping analysis indicated that *mlh3* point mutants displayed genome-wide increases in NCO events that were not observed in wild-type or *mlh3*Δ. Consistent with classical tetrad analysis, *mlh3-32* and *mlh3-D523N* displayed CO values similar to *mlh3Δ*. Importantly, median gene conversion tract lengths associated with COs and NCOs were longer in *mlh3Δ* compared to wild-type, *mlh3-23, mlh3-32* and *mlh3-D523N*. There are multiple possible explanations for these phenotypes, but one possibility is that longer gene conversion tract lengths arise in *mlh3Δ* if the two Holliday junctions present in dHJ intermediates are separated by a longer distance as the result of entry into a pathway that uses a different processing and resolution mechanism (see Discussion). We did not obtain any evidence that the number of DSBs increased in *mlh3* point mutants; such an increase would have provided a simple explanation for why an increase in NCO events was observed. Together, these data provide evidence for *mlh3* separation of function mutants altering the resolution of meiotic recombination intermediates in steps that appear distinct from Mlh1-Mlh3 endonuclease function (see Discussion).

#### i. *mlh3-23, mlh3-32,* and *mlh3-D523N* display distinct CO phenotypes

The average number of COs in *mlh3-23* (92) was similar to that seen in wild-type (93 COs; t test, *P =* 0.31, Table 3; Fig 7B). The *mlh3-32* mutant showed a significant reduction (74 average COs; t-test, *P=*0.0069), with a value comparable to *mlh3Δ* (64 COs, t-test, *P=*0.098). This finding provides additional confirmation that residues K414 and K416 in the Mlh3 linker domain are essential for its meiotic function. Genome wide average CO counts in *mlh3-D523N* (67) were indistinguishable from *mlh3Δ* (64, t-test, *P* = 0.12), consistent with genetic data from a previous single locus study [5]. Analysis of average CO counts per chromosome showed that *mlh3-32* has an intermediate slope between wild-type and *mlh3Δ* (S3A Fig). In addition, *mlh3-32* and *mlh3-D523N* showed significantly reduced COs on medium and large chromosomes (except chromosome I; S3B Fig) like *mlh3Δ* [70]. The CO distribution of *mlh3-23* was similar to wild-type (S3A Fig; S3B Fig).

In wild-type, CO density (cM/kb) varies inversely with chromosome size [71]. All *mlh3* mutants, *mlh3-23* (r = −0.72, p = 0.0015), *mlh3-32* (r = −0.63, p = 0.01), *mlh3-D523N* (r = −0.59, p = 0.02 and *mlh3Δ* (r = -0.76, p = 0.0006) showed a significant negative correlation of CO density (CO/kb) with chromosome size that was similar to wild-type (r = -0.77, p = 5.0 x 10^-4^; Fig 7C).

The median gene conversion tract lengths (S4 Fig) were longer for events associated with crossing over in *mlh3*Δ (2.42 kb) compared to *MLH3* (1.96 kb), *mlh3-23* (1.85 kb), *mlh3-32* (1.80 kb) and *mlh3-D523N* (1.90 kb). This suggests that although *mlh3-32* and *mlh3-D523N* display a decrease in the number of COs that was similar to *mlh3*Δ, COs in *mlh3*Δ may be facilitated through a different mechanism or pathway (see Discussion).

#### ii. Genome-wide increase in non-crossovers in *mlh3-23, mlh3-32* and *mlh3-D523N* mutants

The average number of NCOs in *mlh3-23* (71), *mlh3-32* (68) and *mlh3-D523N* (64) was significantly increased compared to wild-type (46; t-test, *P=*0.00027, 0.0004, and 0.000053, respectively) and *mlh3*Δ (50; t-test, *P=*0.00097, 0.0032, and 0.003, respectively- Table 4; Fig 7B; S3C Fig). In the *mlh3-23* and *mlh3-32* mutants, statistically significant increases in NCOs were observed across small (VI, III), medium (V, X, XIV, II, XIII) and large chromosomes (XII, VII, XV, IV) (S3D Fig). NCO density (NCO/kb) showed significant negative correlation with chromosome size in only wild-type (r = −0.74, p = 0.001) and *mlh3*Δ (r = −0.59, p = 0.02). However, CO plus NCO density showed a significant negative correlation with respect to chromosome size for *mlh3-23* (r = -0.51, p = 0.04), *mlh3-32* (r = -0.67, p = 0.005) and *mlh3-D523N* (r = -0.67, p = 0.004) mutants, suggesting that like *mlh3*Δ, DSB densities are not altered in *mlh3-23*, *mlh3-32* and *mlh3-D523N* mutants.

In addition, the median gene conversion tract lengths were longer for events associated with NCOs in *mlh3Δ* (1.74 kb) relative to *mlh3-23* (1.28 kb), *mlh3-32* (1.17 kb), *mlh3-D523N* (1.35 kb) and *MLH3* (1.50 kb) (S4 Fig; S8 Table). The longer gene conversion tract lengths associated with COs and NCOs in *mlh3Δ* compared to wild-type and the *mlh3* mutants provide support for the idea that Mlh1-mlh3 complexes alter the resolution of meiotic recombination intermediates in steps that appear distinct from Mlh1-Mlh3 endonuclease function (see Discussion).

One explanation for the increase in NCO events seen in *mlh3-23, mlh3-32 and mlh3-D523N* mutants in the genome wide recombination analysis is that these mutants experience meiotic progression delays that result in the continued accumulation of NCOs, possibly through increased DSB formation. Increases in NCO events and a meiotic delay were observed in *ndt80* and the ZMM *zip1*, *zip3* and *msh5* mutants as a result of impeding feedback circuits that inhibit DSB formation [27, 30, 50, 57, 72]. To test this possibility we examined meiotic progression in *MLH3, mlh3Δ, mlh3-32, mlh3-23* and *mlh3-D523N* SK1 strains by measuring the completion of the first meiotic division. This would be difficult to do in S288c/YJM789 strains because they do not show the highly synchronous and efficient meiotic progression profile seen in SK1. As shown in S5 Fig, *MLH3, mlh3Δ, mlh3-32, mlh3-23,* and *mlh3-D523N* mutants showed similar kinetics for completion of at least the first meiotic division (MI+MII), suggesting that the increase in NCO events in *mlh3-32*, *mlh3-23*, and *mlh3-D523N* cannot simply be explained due to a meiotic progression delay. Furthermore, as shown in Fig 7C, the CO plus NCO density data showed a significant negative correlation with respect to chromosome size for *MLH3*, *mlh3-23*, *mlh3-32,* and *mlh3-D523N*, suggesting that DSB densities are unlikely to be altered in the *mlh3* mutants analyzed.

#### iii. Non-exchange chromosome frequencies in mlh3-32 and mlh3-D523N are similar to mlh3Δ

At least one non-exchange chromosome was observed in 43% and 40% of four viable spore tetrads in *mlh3-32* and *mlh3-D523N*, respectively. This was comparable to that seen in *mlh3Δ* (47%; Fig 8A). The non-exchange events in *mlh3-32* and *mlh3-D523N* were observed on small (I, III) and medium size (V, X) chromosomes (S6 Fig). These observations are consistent with crossover defects causing non-exchange events predominantly on the smaller chromosomes as observed previously with *mlh3Δ* and *msh4-R676W* [64, 70]. Lastly, the *mlh3-23* mutant did not show non-exchange chromosomes, consistent with a wild-type number of COs in the S288c/YJM789 hybrid.

**Fig 8.**
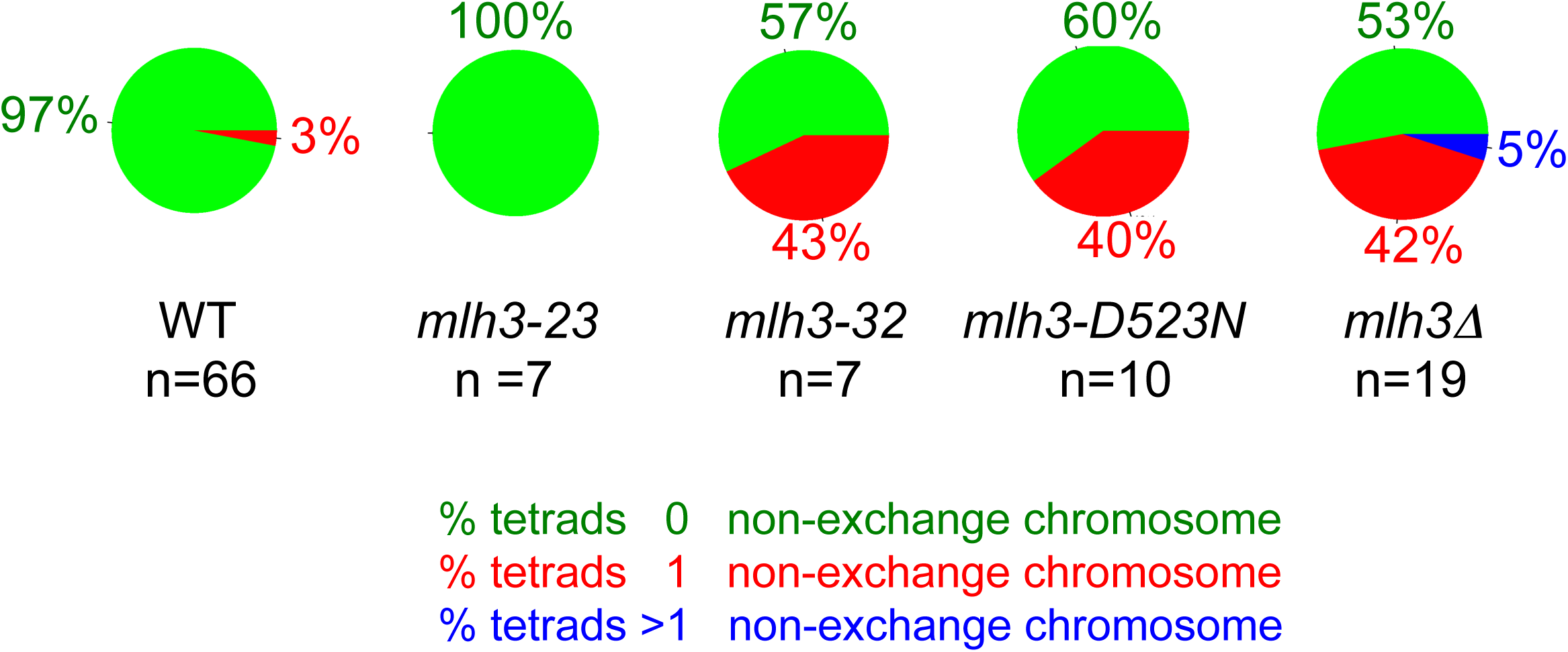
Non-exchange chromosome frequencies in *mlh3-32* and *mlh3-D523N* are similar to *mlh3*Δ and *mlh3-D523N*. Distribution of non-exchange events for wild-type (wt), *mlh3-23, mlh3-32, mlh3-D523N* and *mlh3Δ* in the S288c/YJM789 hybrid background. The percentage of tetrads with zero, one, or more than one non-exchange chromosomes are shown.

## Discussion

We performed a structure-function analysis of Mlh3, a factor that acts in both MMR and meiotic crossing over. This work was pursued because little is known about how Mlh1-Mlh3 acts as a meiotic endonuclease. This is due in part to Mlh1-Mlh3 sharing little in common with the well-characterized structure-selective endonucleases (e.g. Mus81-Mms4, Slx1-Slx4, and Yen1) in terms of homology and intrinsic behavior *in vitro* (reviewed in [51]). Obtaining new mechanistic insights has been complicated by the fact that Mlh1-Mlh3 can bind to model HJ substrates, but cannot cleave them, and by genetic studies suggesting that Mlh1-Mlh3 acts in concert with other pro-CO factors [16, 17, 51]. The identities of these factors are for the most part known, though it is not understood how they contribute to Mlh1-Mlh3’s ability to nick DNA in the directed manner required to generate COs.

Our analysis of two separation-of-function alleles, *mlh3-32* (MMR^+^, CO^-^) and *mlh3-45* (MMR^-^, CO^+^), suggests that protein-protein interactions are critical for directing Mlh1-Mlh3 endonuclease activity (Table 2). Mlh1-Mlh3 has been shown genetically to act downstream of Msh4-Msh5 [40, 41, 44, 48]; this order of events is analogous to steps in DNA MMR where MLH acts following MSH recognition [14, 73]. As outlined in the introduction, Msh4-Msh5, STR, Exo1 (independent of its enzymatic activity) and Zip3 have been classified as pro-CO factors, and have all been shown to interact with one another and/or with Mlh1-Mlh3 (reviewed in [51]). Our biochemical studies are consistent with Mlh1-mlh3-45 having interaction defects that prevent its endonuclease activity from being stimulated by Msh2-Msh3 in MMR. During meiotic CO resolution, we hypothesize that additional factors act in concert to strengthen a possibly weakened Msh4-Msh5-Mlh1-mlh3-45 interaction. This model also helps explain why we identified several MMR^-^ CO^+^ *mlh3* mutants (*mlh3-42* and *-54*) in which the mutant mlh3 protein fails to interact with Mlh1. For the Mlh1-mlh3-32 complex, the MSH interaction and enhancement is retained, but interaction with other critical meiotic factors is likely lost, possibly resulting in an unstable complex that cannot resolve dHJs.

In MMR the asymmetric loading of PCNA by the RFC complex is thought to direct the endonuclease activity of MutLα (Mlh1-Pms1 in *S. cerevisiae*, MLH1-PMS2 in humans) to act in strand-specific repair [74]. Additional studies suggest that specific protein-protein interactions influence and activate MLH endonuclease activity, and direct nicking to a specific location. For example, *in vitro* studies performed with yeast proteins showed that RFC-loaded PCNA can activate Mlh1-Pms1 but not Mlh1-Mlh3 endonuclease on circular plasmids (Mlh3 lacks a PCNA binding motif present in Pms1), and Msh2-Msh3 can activate the endonuclease activity of Mlh1-Mlh3, but not Mlh1-Pms1 [15-17]. Our finding that endonuclease active site residues are highly conserved between Mlh1-Mlh3 and Mlh1-Pms1, which has no role in meiotic crossing over, suggests that the different functions of the two complexes are a result of the different protein-protein interactions.

Data from the Crouse lab suggest that Mlh1-Mlh3 acts in conjunction with Mlh1-Pms1 in Msh2-Msh3 dependent MMR [4]. This observation helps address how Mlh1-Mlh3 is involved in MMR in the absence of a PCNA interaction. Mlh1-Mlh3 is likely recruited and activated by Msh2-Msh3 but also forms a complex with Mlh1-Pms1, which can be directed by PCNA to promote efficient repair. If dimerization between Mlh3 and Mlh1 is weakened, the ability to be recruited by Msh2-Msh3 and interact with Mlh1-Pms1 is likely inhibited, creating a defect in MMR. For meiotic crossing over, a relatively slow process compared to DNA MMR at the replication fork, we suggest that a weakened dimer can be compensated for by interactions with other meiotic factors (e.g. Msh4-Msh5, Exo1 and STR). Thus our work provides further motivation to examine Mlh1-Mlh3 activity on recombination substrates in the presence of pro-CO factors.

### Does Mlh1-Mlh3 have a regulatory role in meiotic pathway choice?

Current meiotic DSB repair models postulate an enzymatic role for Mlh1-Mlh3 in the class I CO pathway after DSB intermediates have been captured and stabilized by the ZMM proteins [8, 30, 50]. In these models DSB intermediates that escape capture by ZMM proteins are resolved into class II COs or NCOs by structure selective nucleases. NCOs can also arise from the action of the STR complex through synthesis dependent strand annealing (SDSA; [30, 50]; Fig 1).

We observed a genome-wide increase in NCOs in *mlh3-23, mlh3-32*, and *mlh3-D523N* that was not seen in *mlh3Δ* mutants or wild-type. In addition, an increase in tract lengths for gene conversions associated with COs and NCOs was observed in *mlh3*Δ compared to wild-type, *mlh3-23, mlh3-32* and *mlh3-D523N* (S4 Fig; S8 Table).

One explanation for the above observations is that in the absence of Mlh1-Mlh3, DSB intermediates are readily available for processing by class II pathway SSNs. In this scenario, Mlh1-Mlh3, in concert with the ZMM proteins, protect recombination intermediates from Sgs1, and thus limit heteroduplex extension. Consistent with this, Zip3 has been shown to limit gene conversion tract lengths by limiting heteroduplex extension driven by Sgs1 [47]. Also, *mlh3Δ, zip3Δ*, and *msh4Δ* show increased CO gene conversion tract lengths that were not seen in the *mlh3* separation of function mutants, suggesting a possible commitment to resolution involving the ZMM proteins and endonuclease-independent functions of Mlh1-Mlh3 [23, 47, 64, 67].

The data presented above can be explained by Mlh1-Mlh3 having an early structural role that is active in *mlh3-23*, *mlh3-32*, and *mlh3-D523N* mutants, resulting in an increase in NCO events due to the loss of biased resolution of dHJs into COs that is a hallmark of the ZMM pathway. Defects in subsequent steps could arise from altered interactions between mutant Mlh1-Mlh3 complexes and pro-CO ZMM factors, permitting structure specific nucleases to resolve dHJs into COs and NCOs (Fig 1). Recent studies suggested that some meiotic factors have earlier roles than first hypothesized; for example, Thacker et al. [75] identified a feedback pathway for the ZMM proteins Zip1, Zip3 and Msh5 that regulates DSB formation in meiosis. At the time this was considered surprising because the ZMM proteins were thought to act exclusively after DSB formation. In support of an early structural role for Mlh1-Mlh3 we found that Sgs1 overexpression decreased the spore viability of *mlh3Δ* strains but not *MLH3* or *mlh3-D523N* strains; we also found that Sgs1 overexpression modestly increased the spore viability of *mlh3-32* mutants (S2 Fig). Interestingly, the spore viability pattern seen in *mlh3Δ* strains overexpressing Sgs1 is consistent with a Meiosis I segregation defect, which might be expected if COs which do not display interference are produced through non-ZMM pathways.

Finally, recent work from Duroc et al. [76] provided evidence that another MLH complex, Mlh1-Mlh2, acts to limit the extent of meiotic gene conversion. In their studies they found that gene conversion tract lengths associated or not associated with COs increased from ~1 kb in wild-type to ~2 kb in *mlh2Δ*. We observed more subtle increases in gene conversion tract length when comparing *mlh3Δ* (1.74 kb for NCOs, 2.42 kb for COs) to wild-type (1.50 kb for NCO, 1.96 kb for COs). These differences could illustrate unique roles for Mlh1-Mlh2 and Mlh1-Mlh3 in regulating meiotic outcomes; however, experiments need to be performed in the same strain background to argue for this idea. Regardless, the fact that Mlh1-Mlh2 does not display endonuclease activity supports the idea that MLH proteins can play structural roles in regulating meiotic recombination outcomes.

It is equally plausible and perhaps simpler that the mutant Mlh1-mlh3 complexes analyzed here display a pathogenic behavior that prevents alternative dHJ resolution activities following ZMM entry. dHJ resolution by Mlh1-Mlh3 is thought to occur when the synaptonemal complex breaks down [9, 77]. If Mlh1-Mlh3 is absent at this time one could imagine that dHJs become susceptible to the actions of the STR complex, resulting in the unwinding and the convergent migration of the two HJs until a single pair of crossing strands in a hemicatenane can be removed by the topoisomerase [30,31]. However, if defective Mlh1-mlh3 complexes remain bound to dHJs and prevent their dissolution by STR, SSNs or other resolvases could resolve dHJs into class II events (Fig 1). This can also explain the longer gene conversion tract lengths associated with COs and NCOs observed in *mlh3Δ* compared to wild-type and the *mlh3* mutants. Physical assays (e.g. two-dimensional electrophoresis) that temporally measure recombination intermediates in meiosis will likely be useful to test this idea (e.g. [8]).

Lastly, it is possible is that delays in meiotic progression in *mlh3* mutants result in the accumulation of NCO events as the result of increased DSB formation [72]. However, we did not observe such delays in any of the *mlh3* mutant backgrounds (S5 Fig). Also, an analysis of the density of CO and NCO events in our genome-wide recombination events suggest that DSB densities were not altered in *mlh3-32*, *mlh3-23*, and *mlh3-D523N* mutants. This is further supported by a decrease in CO:NCO ratios from 2.0 in the wild-type background to 1.3, 1.1, and 1.0 in *mlh3-23*, *mlh3-32*, and *mlh3-D523N* respectively, indicating that the total number of events does not change significantly (from Table 3). These observations suggest that the additional NCO events seen in the *mlh3* mutants did not result from increased DSB formation.

### Mlh3’s linker arm is critical for its meiotic function

MLH proteins act as dimers and contain long unstructured linkers that connect the N- and C-terminal domains of each subunit. These linkers vary in length and are resistant to amino acid substitutions [55]. Previous work showed that the Mlh1-Pms1 heterodimer undergoes large global conformational changes in an ATP binding and hydrolysis cycle [78]. In this cycle the linkers act as arms that can switch between extended and condensed states. These conformational changes are hypothesized to be important to expose different domains of the heterodimer for new protein-protein or protein-DNA interactions in addition to mediating the timing of these interactions [78], and have also been implicated in *B. subtilis* MutL for “licensing” its latent endonuclease activity [60]. In addition, a series of truncation mutants in Mlh1-Pms1 indicate that the Pms1 linker arm appears more important than the Mlh1 linker arm for DNA binding [79]. Extending these ideas to Mlh1-Mlh3, it is interesting to note that the MMR^+^,CO^-^ *mlh3-32* allele maps to the unstructured linker, suggesting that this domain is particularly important in crossing over (Fig 3C), possibly facilitating interactions with CO promoting factors that in turn direct and position Mlh3’s endonuclease activity on recombination substrates. It is important to note that Claeys Bouuaert and Keeney [80] identified mutations in the MLH3 linker domain based on a biochemical analysis of Mlh1-Mlh3 that overlap with residues mutated in the *mlh3-32* allele. Interestingly, the mutations that they identified also conferred a greater defect in crossing over than in DNA mismatch repair, consistent with our analysis of *mlh3-32*. In addition, they found that mutations within and near the *mlh3-32* allele compromised DNA binding activity of Mlh1-Mlh3, suggesting that DNA binding within the linker region may be important for meiotic functions, though we did not detect any apparent defect in the endonuclease activity of Mlh1-mlh3-32.

Alanine-scan mutageneses of Mlh1 [55] and Mlh3 have provided us with additional information regarding the unstructured linkers in Mlh proteins. Previously we used protein structure prediction and molecular analyses to map the Mlh1 unstructured linker to amino acids 336 to 480 [79]; a similar analysis mapped the Mlh3 unstructured linker to amino acids 373 to 490 [16]. As in the analysis of the Mlh3 random coil, few mutations were identified in the Mlh1 unstructured linker that conferred defects in MMR and crossing over. For example in Mlh1, no mutations were identified between amino acids 427 and 490 that conferred mutator phenotypes. However, similar to results seen for Mlh3 (Fig 3C), mutations were identified just before the unstructured linker in Mlh1 (253-312) that conferred strong mutator phenotypes [55]. Curiously, the corresponding region in MutL contains residues that have been linked through crystallographic analysis to DNA binding [81], suggesting that the organization of the DNA binding and unstructured linker domains in the MLH proteins is conserved. Finally, in both Mlh1 and Mlh3, a localized set of mutations within the center of the unstructured linker (390-403 in Mlh1, 414-416 in Mlh3) affect function, suggesting that this specific region is likely to play an important function beyond serving as a random coil.

### Closing thoughts

Mlh1-Mlh3 appears to be acting in CO resolution through a novel mechanism distinct from known structure-selective endonucleases. Mlh1-Mlh3 does not share conservation with the known endonuclease superfamilies (XPF, URI-YIG, Rad2/XPG), and does not appear capable of resolving model HJ substrates [51]. As mentioned previously, dHJ resolution by Mlh1-Mlh3 results in only CO products whereas the interference-independent CO pathway, which is dependent on Mus81-Mms4, resolves dHJs into a mixture of CO and NCO products [8]. Thus, Mlh1-Mlh3’s distinct activity suggests that its nicking is positioned by pro-CO factors such as Msh4-Msh5, Zip3, the STR complex, and Exo1. Such factors are likely to orient Mlh1-Mlh3 to promote asymmetric cleavage of dHJs in a highly regulated and coordinated manner. Thus our work provides further motivation to examine Mlh1-Mlh3 activity on recombination substrates in the presence of pro-CO factors.

Polymorphisms in human *MLH3* genes have been associated with male and female infertility [82-84], and errors in meiotic chromosome segregation are considered a leading cause of spontaneous miscarriages and birth defects [13]. It is interesting to note that the *mlh3-23* mutation, which only weakly affected crossing over, conferred an alteration in meiotic recombination outcomes that was similar that seen in *mlh3* mutants that conferred more severe defects (Fig 7). This observation suggests that some polymorphisms in meiotic recombination genes could have more severe defects in human fertility than expected.

## Methods

### Media

*S. cerevisiae* SK1, S288c, and YJM789 strains were grown on either yeast extract-peptone-dextrose (YPD) or minimal complete media at 30°C [85]. For selection purposes, minimal dropout media lacking uracil was used when needed. Geneticin (Invitrogen, San Diego) and nourseothricin (Werner BioAgents, Germany) were added to media when required at recommended concentrations [86, 87]. Cells were sporulated as described by Argueso et al. [7].

### Site-directed mutagenesis of *MLH3*

60 *mlh3* alleles were constructed, resulting in the mutagenesis of 139 amino acids in the 715 amino acid Mlh3 polypeptide (S1 Table). The single-step integration vector (pEAI254), containing the SK1 *MLH3* gene with a *KANMX4* selectable marker inserted 40 bp downstream of the stop codon [5], was used as a template to create plasmids bearing the *mlh3* mutant alleles via QuickChange site directed mutagenesis (Stratagene, La Jolla, CA). *mlh3-60*, in which the last 11 residues of Pms1 (DWSSFSKDYEI) were inserted before the *MLH3* stop codon, was also made by QuickChange. Mutations were confirmed by sequencing the entire open reading frame (Sanger method), as well as 70 bp upstream and 150 bp downstream. Primer sequences used to make and sequence these variants are available upon request.

### Mlh3 homology model

The amino acid sequence of *S. cerevisiae* Mlh3 (YJM789) was used to construct a homology model from HHpred (http://toolkit.tuebingen.mpg.de/hhpred) and Modeller software. PyMOL was used for imaging.

### Construction of strains to measure meiotic crossing over and MMR

The SK1 strain EAY3255 (S1 Table) was constructed to allow for the simultaneous analysis of *mlh3* MMR and meiotic crossing over phenotypes. It carries a spore autonomous fluorescent protein marker (RFP) on chromosome VIII to monitor chromosome behavior (crossing over and non-disjunction; [57]) as well as the *lys2::InsE-A*_*14*_ cassette to measure reversion to Lys^+^ [56]. pEAI254 and mutant derivatives described above and in S3 Table were digested with *Bam*HI and *Sal*I and introduced into EAY3255 by gene replacement using the lithium acetate transformation method as described in Gietz et al. [88]. At least two independent transformants for each genotype (verified by sequencing) were made resulting in a total of 120 haploid strains bearing the *mlh3* variants described in this study (S1 Table). These haploid strains were used to measure the effect of *mlh3* mutations on reversion rate and were mated to EAY3486, an *mlh3*Δ strain containing the CFP marker, resulting in diploid strains suitable for analysis of crossing over (S2 Table). Diploids were selected on media lacking the appropriate nutrients and maintained as stable strains. Meiosis was induced upon growing the diploid strains on sporulation media as described in Argueso et al. [7]. Wild-type strains carrying the fluorescent protein markers used to make the above test strains were a gift from the Keeney lab.

### Lys^+^ reversion assays

The haploid strains described above were analyzed for reversion to Lys^+^ as described in Tran et al. [56]. At least 10 independent cultures were analyzed for each mutant allele alongside wild-type or *mlh3*Δ controls. Analyses were performed for two independent transformants per allele. Reversion rates were measured as described [89, 90], and each median rate was normalized to the wild-type median rate (1X) to calculate fold increase. Alleles were classified into a wild-type, intermediate, or null phenotype based on the 95% confidence intervals which were determined as described [91].

### Spore autonomous fluorescent protein expression to measure percent tetratype

Diploids in the EAY3255/EAY3486 background described above (S2 Table) were sporulated on media described in Argueso et al. [7]. Spores were treated with 0.5% NP40 and briefly sonicated before analysis using the Zeiss AxioImager.M2 [57]. At least 250 tetrads for each *mlh3* allele were counted to determine the % tetratype. Two independent transformants were measured per allele. A statistically significant difference (p<0.01) from wild-type and *mlh3Δ* controls based on χ^2^ analysis was used to classify each allele as exhibiting a wild-type, intermediate, or null phenotype.

### Meiotic time courses

Meiotic time course were performed as described in Sonntag Brown et al. [36] for the diploid strains EAY3252/EAY3486 (*MLH3*), EAY3255/EAY3486 (*mlh3Δ*), EAY3534-35/EAY3486 (*mlh3-23*), EAY3552-53/EAY3486 (*mlh3-32*), and EAY3819-20/EAY3486 (*mlh3-D523N*; S2 Table). Strains in single time courses were grown in the same batch of media under identical conditions. Aliquots of cells at specific time points were stained with DAPI to determine the percentage of cell that completed the first meiotic division (cells in which 2, 3, or 4 nuclei were observed by DAPI staining, presented as MI+MII). Cells were visualized using a Zeiss Axio Imager M2 microscope equipped with a DAPI filter. At least 150 cells were counted for each time point. Two independent transformants were analyzed per allele.

### Yeast two-hybrid analysis

The L40 strain [92] was co-transformed with bait and target vectors. Residues 481-715 of the Mlh3 C-terminus were PCR amplified from pEAI254 (SK1 *MLH3* described above) and mutant derivatives, and then sub-cloned into the target vector pEAM98 (S288C *MLH3*). pEAM98 contains a fusion between the GAL4 activation domain in pGAD10 and residues 481-715 of the Mlh3 C-terminus [5, 35]. The resulting SK1 derived target vectors were confirmed by sequencing (Sanger method). The bait vector used was (pBTM-Mlh1) as described in Nishant et al. [5]. Expression of the *LACZ* reporter gene was determined by the ortho-nitrophenyl-β-D-galactopyranoside (ONPG) assay as described in [93].

### Purification of Mlh1-Mlh3 and mutant complexes from baculovirus- infected Sf9 cells

Mlh1-Mlh3 and Mlh1-mlh3 mutant derivatives were purified from Sf9 cells infected with Bac-to-Bac baculovirus expression system using pFastBacDual constructs [16]. Mutant Mlh1-mlh3 complexes were purified using the same protocol developed to purify wild-type Mlh1-Mlh3. This involved the use of successive nickel-nitroloacetic acid-agarose (Qiagen) and heparin sepharose (GE Healthcare) column purifications. Mlh1-Mlh3 and mutant derivative yields were ~150 μg per 5 × 10^8^ cells; aliquots from the final heparin purification were frozen in liquid N_2_ and stored at −80°C. Protein concentrations were determined by Bradford assay [94] using BSA standard. The *mlh3-6*, *mlh3-32* and *mlh3-45* mutations were introduced into pEAE358 (pPH-His_10_-MLH3-HA pFastBacDual construct; Rogacheva et al. [16]) by Quick Change (Stratagene). *His*_*10*_*-mlh3-HA* fragments were individually subcloned by restriction digestion into pEAE348 to form pFastBacDual constructs pEAE382 (Mlh1-mlh3-6), pEAE383 (Mlh1-mlh3-32) and pEAE384 (Mlh1-mlh3-45), in which the *MLH1-FLAG* gene is downstream of the p10 promoter and the *His*_*10*_ *-mlh3-HA* gene is downstream of the pPH promoter. The sequence of the restriction fragments inserted into pEAE348 were confirmed by DNA sequencing (Cornell Biotechnology Resource Center). Msh2-Msh3 was purified as described previously [95].

### Endonuclease assay on supercoiled plasmid DNA and ATPase assay

Mlh1-Mlh3 nicking activity was assayed on supercoiled pBR322 or pUC18 (Thermo Scientific). DNA (2.2 nM) was incubated in 20 μl reactions containing indicated amounts of Mlh1-Mlh3 and Msh2-Msh3 [95] in 20 mM HEPES-KOH pH 7.5, 20 mM KCl, 0.2 mg/ml BSA, 1% glycerol, and 1 mM MgCl_2_ for 1 h at 37°C. Reactions were quenched by incubation for 20 min at 37°C with 0.1% SDS, 14 mM EDTA, and 0.1 mg/ml proteinase K (New England Biolabs) (final concentrations). Samples were resolved by 1% agarose gel with 0.1 μg/ml ethidium bromide run in 1X TAE buffer for 50 min at 95 V. All quantifications were performed using GelEval (FrogDance Software, v1.37). The amount of nicked product was quantified as a fraction of the total starting substrate in independent experiments. *bkg* indicates that amount of nicked product was not above background levels established by negative controls. ATPase assays were performed as described [16].

### Genetic map distance analysis

Diploids from the SK1 congenic strain background EAY1112/1108 [5-7] were used for genetic map distance analyses. EAY1112/2413 (*MLH3/mlh3Δ::NATMX*) and EAY1848/2413 (*mlh3Δ::KANMX/mlh3Δ::NATMX*) were used as wild-type and null controls respectively (S2 Table). *mlh3* alleles of interest were integrated into EAY3712 (same as EAY1112 but *mlh3Δ::URA3*) using standard techniques [88]. The resulting haploid strains (EAY3713-EAY3724) were mated to EAY2413 (*mlh3Δ::NATMX*) giving rise to diploids carrying markers suitable for genetic distance measurements (S2 Table). Two independent transformants were analyzed per allele. Diploids were selected on media lacking the appropriate nutrients and maintained as stable strains. Diploids were sporulated as described [7]. Tetrads were dissected on synthetic complete media and germinated at 30°C after an incubation of 2-3 days. Spore clones were then replica-plated on various selective media to be scored after 1 day of incubation at 30°C. Chromosome behavior was analyzed using the recombination analysis software RANA to measure genetic map distances and spore viabilities [7]. Genetic map distances ± SE were calculated using the formula of Perkins [96] through the Stahl Laboratory Online Tools portal (http://molbio.uoregon.edu/~fstahl/).

### Construction of strains for whole genome sequencing

Strain genotypes are shown in S1 Table. SK1-*MLH1 and MLH3* alleles were introduced into wild-type YJM789 and S288c *mlh3Δ::natMX4*, respectively, using plasmids pEAA214 and pEAI254. The SK1 *MLH1* specific SNPs were confirmed by Sanger sequencing. The *mlh3-23::kanMX4, mlh3-32::kanMX4*, and *mlh3-D523N::kanMX4* mutations were introduced using plasmid pEAI347, pEAI356, and pEAI252 respectively in a S288c *mlh3Δ::natMX4* background. The S288c *mlh1Δ::hphMX4* and YJM789 *mlh3Δ::kanMX4* strains were made using deletion constructs amplified by PCR.

### Genome wide mapping of meiotic recombination events in the S288c/YJM789 hybrid

Genomic DNA was extracted from spore colonies of four viable spore tetrads of the *mlh3* mutants as described previously [64]. Whole genome sequencing on the Illumina Hi-Seq 2500 platform was performed at Fasteris, Switzerland. Raw sequence reads were processed and SNPs genotyped as described in Chakraborty et al. [70]. Analysis of recombination events, interference was performed using the CrossOver program (v6.3) in the ReCombine suite of programs (v2.1; [66]). Parameters for the CrossOver program were set as described in Krishnaprasad et al. [64]). Custom R scripts were used to generate the segregation file (input file for the CrossOver program), plots and to perform statistical tests. The raw recombination data files and the custom R scripts are available online at the Dryad digital repository (http://datadryad.org; doi:10.5061/dryad.bb702). Sequence data are available from the National Centre for Biotechnology Information Sequence Read Archive (Accession number SRP096621).

## Overexpression of Sgs1

*SGS1* (native promoter, ORF, and termination sequence) was PCR amplified from SK1 genomic DNA obtained from NKY730 (*MATa/alpha, ura3Δ::hisG/ura3Δ::hisG, leu2::hisG/leu2::hisG, lys2/lys2*) and cloned into the high copy vector pEAO34 (pRS426: 2μ, Amp^R^, *URA3*). The correct DNA sequence was confirmed by Sanger sequencing the entire insert. Primer sequences used to make this construct are available upon request. The high copy vector (with or without the *SGS1* insert) was transformed into stable diploids of EAY3819-20/EAY3486 (*mlh3-D523N/mlh3Δ*), EAY3552-53/EAY3486 (*mlh3-32*/*mlh3Δ*), EAY3252/EAY3486 (*MLH3/mlh3Δ*) and EAY3255/EAY3486 (*mlh3Δ/mlh3Δ*) backgrounds as controls (S2 Table). Meiosis was induced as described in Argueso et al. [7] and vector selection was maintained by growing the diploid strains on minimal media lacking uracil prior to sporulation. In addition, sporulation media lacked uracil. For spore viability measurements, tetrads were dissected on synthetic complete media and germinated at 30°C after an incubation of 2-3 days. Two independent transformants were analyzed per high copy vector. Differences in spore viability were assessed for significance using the χ^2^ test.

## Acknowledgements

We are grateful to Michael Lichten, Nancy Kleckner, Alba Guarne, Duyen Bui, Ujani Chakraborty, and Christopher Furman for fruitful discussions, Scott Keeney for reagents, Nathan Kruer-Zerhusen for help constructing the *S. cerevisiae* Mlh3 homology model, and the Cornell Statistical Consulting Unit for help with the statistical analysis of the *lys2-A*_*14*_ data. We are thank Scott Keeney and Corentin Claeys Bouuaert for insightful comments and for sharing unpublished data.

## Supporting Information

**S1 File. Segregation of SNPs in all 28 tetrads as labeled in S6 Table.** S288c and YJM789 SNPs are shown in red and blue, respectively.

**S1 Fig. Mlh1-mlh3-6 exhibits wild-type endonuclease and ATPase activity.** A. SDS-PAGE analysis of purified Mlh1-Mlh3 and Mlh1-mlh3-6. Coomassie Blue R250-stained 8% Tris-glycine gel. 0.5 μg of each protein is shown. MW = Molecular Weight Standards from top to bottom-116, 97, 66, 45, 31 kD). B, C. Mlh1-Mlh3 and Mlh1-mlh3-6 (18, 37, 70 nM) were incubated with 2.2 nM supercoiled pBR322 DNA, analyzed in agarose gel electrophoresis, and endonuclease activity was quantified (average of 6 independent experiments presented +/-SD) as described in the Experimental Procedures. C. ATPase assays were performed as described in Rogacheva et al. [16], but contained the indicated amounts of Mlh1-Mlh3 and Mlh1-mlh3-6 incubated with 100 μM ^32^P-γ-ATP. Reactions were performed in duplicate for two separate purifications of each, and the average values, +/-SD, are presented.

**S2 Fig. Sgs1 overexpression differentially affects spore viability in *mlh3Δ* vs. *MLH3* and *mlh3-D523N* strains.** A. Distribution of viable spores in tetrads of *MLH3, mlh3Δ*, and *mlh3-D523N* strains containing p*SGS1-2μ*. B. Distribution of viable spores in tetrads of *mlh3-32* strains containing p*SGS1-2μ*. In all plots, the horizontal axis corresponds to the classes of tetrads with 4, 3, 2, 1 and 0 viable spores, and the vertical axis corresponds to the frequency of each class given in percentage. The overall spore viability (SV) and the total number of spores counted (n) are shown.

**S3 Fig. Crossover and non-crossover distribution on chromosomes for wild-type, *mlh3-23, mlh3-32, mlh3-D523N* and *mlh3Δ*.** A. and C. Scatter plot of average crossover (CO) and non-crossover (NCO) counts per chromosome against chromosome size. The equations for the regression lines are: wild-type (CO = 0.0000063 * chr. size + 1.09; NCO = 0.0000032 * chr. size + 0.44), *mlh3-23* (CO = 0.0000063 * chr. size + 1.02; NCO = 0.0000061 * chr. size - 0.16), *mlh3-32* (CO = 0.0000049 * chr. size + 0.92, NCO = 0.0000053 * chr. size + 0.25), *mlh3-D523N* (CO = 0.0000042* chr. Size + 1.05, NCO = 0.0000046 * chr. size + 0.52), *mlh3Δ* (CO = 0.0000041 * chr. size + 0.91; NCO = 0.0000032 * chr. size + 0.69). B. and D. Bar plot of average crossover and non-crossover counts per chromosome. The asterisk symbol (*) marks chromosomes that have significant difference (two tailed t-test for difference in mean; P<0.05) in crossover / non-crossover counts compared to wild-type. Chromosomes are arranged by size from left to right. Error bars are mean ± std. error.

**S4 Fig. Gene conversion tract lengths associated with crossovers and non-crossovers for wild-type, *mlh3-23, mlh3-32, mlh3-D523N* and *mlh3Δ*.** A. Distribution of gene conversion tract lengths (outlier points not shown) associated with crossovers and non-crossovers for wild-type, *mlh3-23, mlh3-32, mlh3-D523N* and *mlh3Δ*. Statistically significant differences are marked by asterisks (*, 0.05 > P > 0.001); (***, 0.0001 > P). B. and C. Breakdown of the number of crossover and non-crossover gene conversion tracts in sizes ranging from 0 to 20 kb in 500 bp intervals. Median tract lengths are shown in dotted lines and in S8 Table.

**S5 Fig. *mlh3* mutants display normal meiotic prophase progression as measured by the completion of the first meiotic division.** Representative time courses showing the completion of the MI division (MI+MII) in *MLH3, mlh3-23, mlh3-32, mlh3-D523N* and *mlh3Δ* strains. Cells with two, three, or four nuclei were counted as having completed MI (MI+MII). All strains for a single time course were grown in the same batch of media under identical conditions. Two independent transformants were measured per allele.

**S6 Fig. The percentage of meioses that show a non-exchange for the indicated chromosome.** Chromosomes are shown with respect to increasing size.

**S1 Table. Yeast strains used in this study.** EAY3252, EAY3255 and derivatives, and EAY3486 are SK1 strains that contain spore-autonomous fluorescent markers described in Thacker et al. [57]. EAY1112 and EAY2413 contain chromosome XV markers described in Argueso et al. [7]. S288c, YJM789 and derivative KTY strains were used for whole genome recombination mapping as described in Mancera et al. [67] and Krishnaprasad et al. [64].

**S2 Table. Diploid strains used to measure % tetratype, spore viability, meiotic progression, genetic map distances and for whole genome recombination mapping.** The indicated haploid strains were mated to form the diploids with the relevant genotype shown.

**S3 Table. Plasmids used in this study.** A. All *MLH3* mutagenesis plasmids are derived from pEAI254, a 7.8 kb *MLH3*_*SK1*_::*KANMX* integrating vector. pEAI254 was mutagenized by QuickChange to create the alleles listed. The DNA sequence of the entire ORF, and 70 bp upstream and 150 bp downstream, were confirmed by DNA sequencing using primers EAO318, EAO319, EAO1778 and EAO321. B. For the two-hybrid analysis, pEAM105 contains the entire *MLH1* gene derived from the SK1 strain inserted immediately after the lexA binding domain in pBTM116. All *GAL4* activating domain-*mlh3* plasmids are derived from pEAM234, which contains DNA sequence encoding SK1 MLH3 amino acids 481 to 715 inserted immediately after the GAL4 activating domain in pGAD424.

**S4 Table. Genetic map distances for *mlh3* separation of function mutants on chromosome XV from single spores and tetrads.** All *mlh3* mutants are isogenic derivatives of EAY1112/EAY2413 (S2 Table; Methods). For single spores, recombination frequencies (recombinant spores/total spores) were multiplied by 100 to yield genetic map distances (cM). For tetrads, genetic distance in centimorgans (cM) was calculated using the RANA software without considering aberrant segregants (Argueso et al. [7]). The Stahl Laboratory Online Tools website (http://molbio.uoregon.edu/~fstahl/) was used to calculate standard error (SE) around the genetic distance for tetrads. n; number of single spores, N; four spore viable tetrads analyzed; Par, parental single spores; Rec, recombinant single spores.

**S5 Table. Sequencing statistics for spores derived from *mlh3-23, mlh3-32, mlh3-D523N* and wild-type S288c/YJM789 hybrid bearing SK1-*MLH1/MLH3* alleles** *mlh3Δ* spore sequencing statistics can be found in Chakraborty et al. [70].

**S6 Table. Crossovers (CO), non-crossovers (NCO) and non-exchange chromosomes (NEC) in tetrads of *mlh3-23, mlh3-32, mlh3-D523N* and wild-type S288c/YJM789 hybrid bearing SK1-*MLH1/MLH3* alleles.** *mlh3Δ* data can be found in Chakraborty et al. [70].

**S7 Table. Average crossovers (CO) and non-crossovers (NCO) per chromosome for WT (wild-type), *mlh3-23, mlh3-32, mlh3-D523N* and *mlh3Δ* mutants.** *Merged data set from [64, 67]. ^#^Data set from Chakraborty et al. [70].

**S8 Table. Segregation frequency of SNP markers and gene conversion tract lengths in *mlh3Δ*, wild type, *mlh3-23, mlh3-32* and *mlh3-D523N* mutants.** The percentage of SNP markers with 2:2, 3:1, 1:3, 4:0 and 0:4 segregation (S288c:YJM789) and the median gene conversion tract lengths in kb for crossovers (CO) and non-crossovers (NCO) are shown. *P*-values show the statistical significance of the difference in median CO and NCO gene conversion tract lengths of wild-type and *mlh3* alleles compared to *mlh3∆*.

